# PDL-1+ Neutrophils mediate susceptibility during endotoxemia in Metabolic Dysfunction–Associated Steatotic Liver Disease

**DOI:** 10.1101/2024.10.16.618651

**Authors:** Cleyson da Cruz Oliveira Barros, Alexandre Kanashiro, Gabriel Victor Lucena da Silva, Paulo Sérgio de Almeida Augusto, Guilherme Cesar Martelossi Cebinelli, Luiz Osório Leiria, Thiago Mattar Cunha, José Carlos Alves Filho, Fernando Queiroz Cunha

**Author notes:** These authors contributed equally to this work (Cleyson da Cruz Oliveira Barros, Alexandre Kanashiro).

## Abstract

Metabolic Dysfunction–Associated Steatotic Liver Disease (MAFLD) is a pathological condition affecting many individuals worldwide. Patients with MAFLD are more susceptible to systemic inflammation, including endotoxemia, which accelerates the progressive liver damage. However, the immunological mechanisms that trigger the hyper-inflammatory responses in individuals with MAFLD remain unknown. In the present study, we reported that short-term HFCD (Choline Deficient High Fat Diet)-fed mice, which did not show significant signs of hepatic damage and inflammation in the first two weeks, are more susceptible to a non-severe sepsis-like systemic inflammation induced by LPS challenge. Mechanistically, endotoxemic mice show an excessive accumulation of NK-producing IFN-γ cells in liver tissue, which trigger the recruitment and polarization of a distinct subset of neutrophils, characterized by high expression of PD-L1 and massive release of TNF-α. Remarkably, genetic inhibition of IFN-γ or pharmacological blockade of PD-L1 effectively modulated the excessive recruitment of these neutrophils to the liver and TNF-α production, thereby preventing hepatic damage and reducing the severity of host mortality. Thus, these results support the design of novel effective strategies to control hyperinflammatory responses in septic HFCD patients and consequently improve their survival.

## 1. INTRODUCTION

Metabolic Dysfunction–Associated Steatotic Liver Disease (MAFLD), previously described as non-alcoholic fatty liver disease (NAFLD), is a metabolic liver dysfunction defined as an abnormal accumulation of fat in the liver without secondary triggers such as alcohol consumption. MAFLD has become a common chronic liver disease in the world population and has a high prevalence varying from simple steatosis to non-alcoholic steatohepatitis, fibrosis, cirrhosis, and hepatocellular carcinoma (1–3). Despite its high prevalence and increasing cause of liver-related mortality worldwide, MAFLD is still underestimated, and there are no approved pharmacological or preventive strategies for these medical conditions (4, 5).

The MAFLD is a highly complex condition that involves a “multi-hit” process, in which different stimuli act together to promote fat accumulation and progressive liver damage (6). Although the pathogenesis of MAFLD is multifactorial, inflammation has now been considered a key factor in the progression of advanced hepatic conditions (7–9). In the presence of obesity and lipotoxicity, the immune system is activated in the liver, with a corresponding degree of inflammation triggered by the “wound healing response”. Since these inflammatory signs persist for a long period, monocytes and neutrophils, as well as T CD4, T CD8, and natural killer (NK), can be recruited to the liver, releasing a massive secretion of pro-inflammatory cytokines, including interferon-gamma (IFN-γ) and tumor necrosis factor (TNF-α) (7, 10). Likewise, in chronic diseases, these cytokines are also present in systemic and uncontrolled inflammatory processes, such as sepsis (11). Sepsis is a syndrome characterized by a life-threatening dysfunction due to complex interactions between pathogens and the host recruitment of immune cells, resulting in a systemic hyper-inflammatory state followed by extensive vital organ damage, including in the liver (12, 13). Sepsis has also been considered the leading cause of death in intensive care units. In endotoxemia, a well-established sepsis model, this massive production of cytokines provokes uncontrolled recruitment of innate immune cells, mainly monocytes and neutrophils, to vital organs, including the liver, where they induce extensive and lethal damage (11, 14, 15). The majority of data regarding endotoxemia pathogenesis has focused only on the interaction between the infection and the host immune system, and only a few studies have addressed one or more clinical conditions that are considered risk factors leading to endotoxemia aggravation, such as senescence and previous immunosuppressive comorbidities (16–18). Thus, understanding how these clinical conditions can modify the host immune response and consequently affect the outcome of sepsis is crucial to detecting the risks and designing novel preventive and therapeutic approaches to control exacerbated inflammatory responses. Recently, some population studies, including large-scale data from cohorts with biopsy-confirmed individuals, have evidenced a higher incidence of severe infections in MAFLD patients, suggesting that previous hepatic conditions can modify the host immune response, leading to sepsis aggravation (19). However, although a relationship between elevated risk of sepsis in MAFLD patients exists, the underlying immunological mechanisms involved in the progression and exacerbation of systemic and/or local (hepatic) inflammation in these individuals have not been identified yet.

To understand the impact of MAFLD on liver damage during endotoxemia, we used a known murine model of MAFLD and later challenged these animals with a moderate dose of lipopolysaccharide (LPS). We demonstrated that endotoxemia exacerbated the hepatic inflammatory response and mortality in Choline Deficient High Fat Diet (HFCD)-fed mice. The LPS-activated hepatic NK cells secrete IFN-γ, which induces massive recruitment and polarization of a specific subpopulation of PD-L1-expressing neutrophils. This subpopulation of neutrophils releases a high concentration of TNF-α, which mediates the liver lesion and increases the mortality of the animals. In fact, biological treatments promoting PD-L1 neutralization, or neutrophil depletion, reduced the TNF-α concentration, the liver inflammatory response and organ pathology, and increased the survival in endotoxemic HFCD fed mice. Similar results were observed with the use of genetic knockout TNF-α receptor mice. Thus, our findings describe the potential mechanism involved in the exacerbation of neutrophil functions in MAFLD mice and suggest a viable approach to the development of individual interventions capable of mitigating the hyperinflammation observed in these pathological conditions.

## 2. RESULTS

### MAFLD mice are more susceptible to LPS-induced systemic inflammation

To study MAFLD as a pre-condition associated with increasing risk of exacerbation of inflammation induced by endotoxemia, we employed a murine model of MAFLD in which the mice were fed for two weeks with a choline-deficient high-fat HFCD diet, a well-known liver disease model (Fig. 1A). Therefore, we adopted a two-week HFCD feeding period, during which no changes in lymphocyte numbers were observed, as longer exposure to the choline-deficient HFCD is known to reduce lymphocyte numbers in mice (20). Mice fed HFCD showed no increase in liver weight and collagen deposition as evidenced by picrosirius staining (Fig. S1A and Fig. S1C). The HFCD mice challenged with LPS (10 mg/kg; intraperitoneally) triggered an elevated susceptibility with an index of mortality of 100% 24 h after the challenge (Fig. 1B). In our experimental condition, this dose of LPS provoked moderate (below 40%, 4 days after LPS challenge) mortality in chow-fed mice (control group) (Fig. 1B). Compared with the control group, the HFCD animals challenged with LPS showed elevated liver damage, evidenced by an increase in plasma alanine aminotransferase (ALT) levels (a known marker of liver injury) and histopathological alteration, characterized by leukocyte infiltration and edema (Fig 1C,1E). Compared with control mice treated with PBS, the HFCD animals also treated with PBS did not show significant signs of hepatic damage and local inflammatory response, evidenced by the absence of leukocyte infiltration (Fig. 1C and 1E). Animals subjected to the HFCD diet showed reduced blood glucose levels with or without LPS challenge (Fig. S1B). Remarkably, compared to the Chow group, HFCD mice exposed to LPS did not show greater changes in other organs commonly affected by endotoxemia, such as the kidneys (Figure 1D), indicating that a pathological inflammatory condition is increased mainly in liver tissue compared to other organs. To assess how MAFLD modulates the immune response in the hepatic microenvironment of human patients, we reanalyzed previously published bulk RNA-seq data from patients with MAFLD to identify potential targets related to the inflammatory process (21). First, we performed analyses of genes differentially expressed in leukocyte samples from patients with MAFLD and healthy controls. We observed that 5240 genes are upregulated, such as *EIF4H*, *TMEM*, *MOB1A*, *SERBP1*, *TM9SF1*, *HINT3*, *ARF3*, *CHTF8*, and *PDIA6*, while 5806 genes, as *FOSB*, *KAT2A*, *SNRNP70*, *CCNL2*, *AMY2B*, *EXD3*, and *EBNA1BP2, are* downregulated. Interestingly, we found that the *CD274* gene (encoding PD-L1) and the *IFNGR2* gene, corresponding to the IFN-γ receptor, are among the upregulated genes (Fig. 2A). Furthermore, after conducting an additional enrichment analysis of the 5240 upregulated genes using Gene Ontology (GO), we detected an increase in several pathways associated with inflammatory processes, such as phagocytosis, leukocyte migration, immune response to bacterial components, and signaling pathways involving IL-1β, TNF-α, and IFN-γ (Fig. 2B).

**Figure 1.**
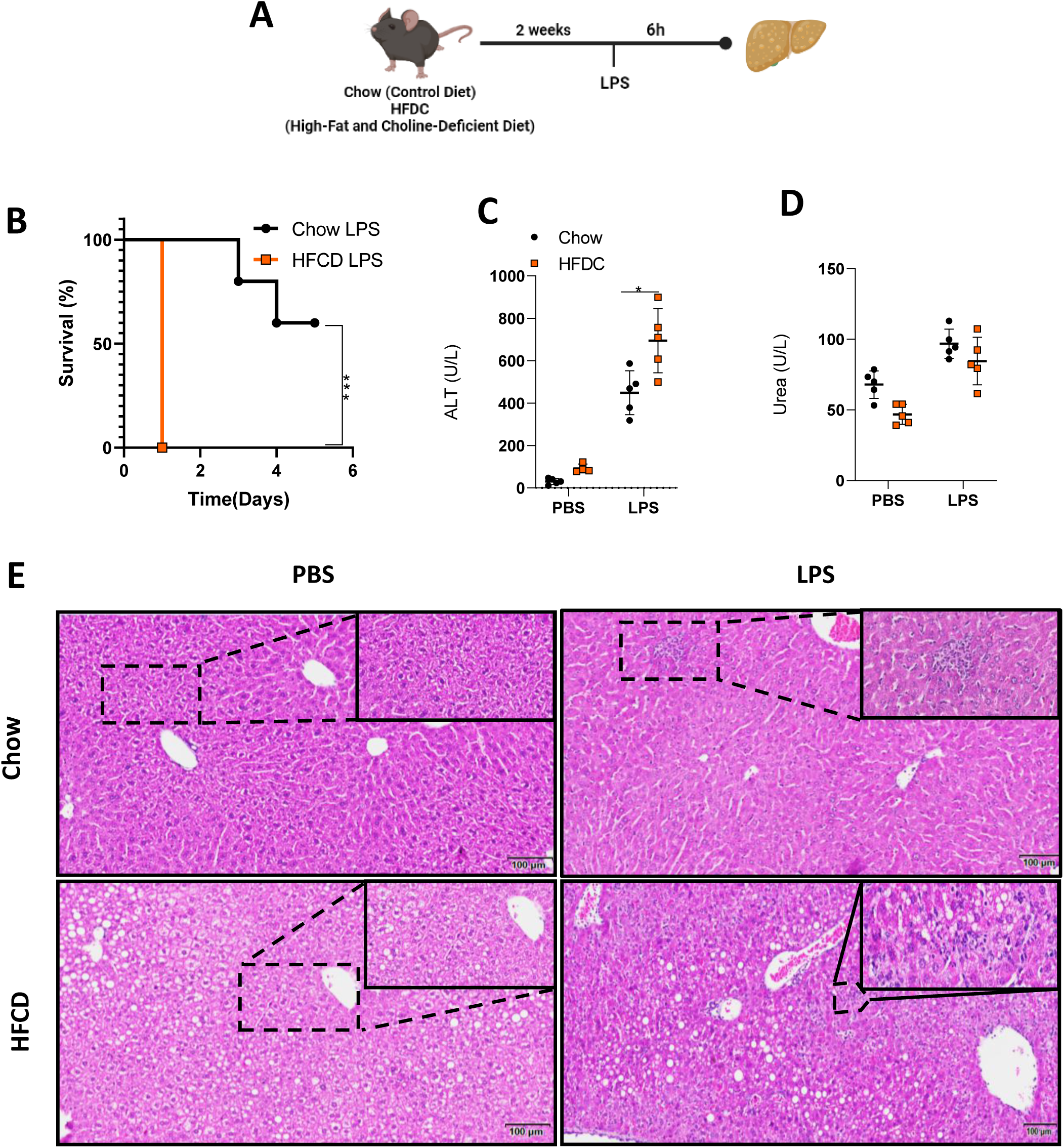
Mice fed with a high-fat and choline-deficient (HFCD) diet develop Metabolic Dysfunction–Associated Steatotic Liver Disease (MAFLD) and are more susceptible to endotoxemia. (A) Schematic illustration of the experimental design. (B) Survival curves of mice with MAFLD (n=5) and Chow diet (controls) (n=5) which were injected intraperitoneally with LPS (10mg/kg) and the survival rates were determined daily for 7 days (C and D) Serum ALT and Urea levels from mice with MAFLD and Chow, 6 hours after LPS (10 mg/kg) and PBS administration (n=5). (E) Effect of LPS inoculation on histopathology (H&E stain) of liver from mice with MAFLD or controls (Chow diet). The results are expressed as the mean ± standard deviation (SD) from one representative experiment. Statistical analysis was performed using one-way ANOVA followed by Tukey’s multiple comparison test. Statistical significance was determined with the following p-values: *p < 0.05, **p < 0.01, ***p < 0.001. Each experiment was independently repeated 2 times to ensure reproducibility of the findings.

**Figure 2.**
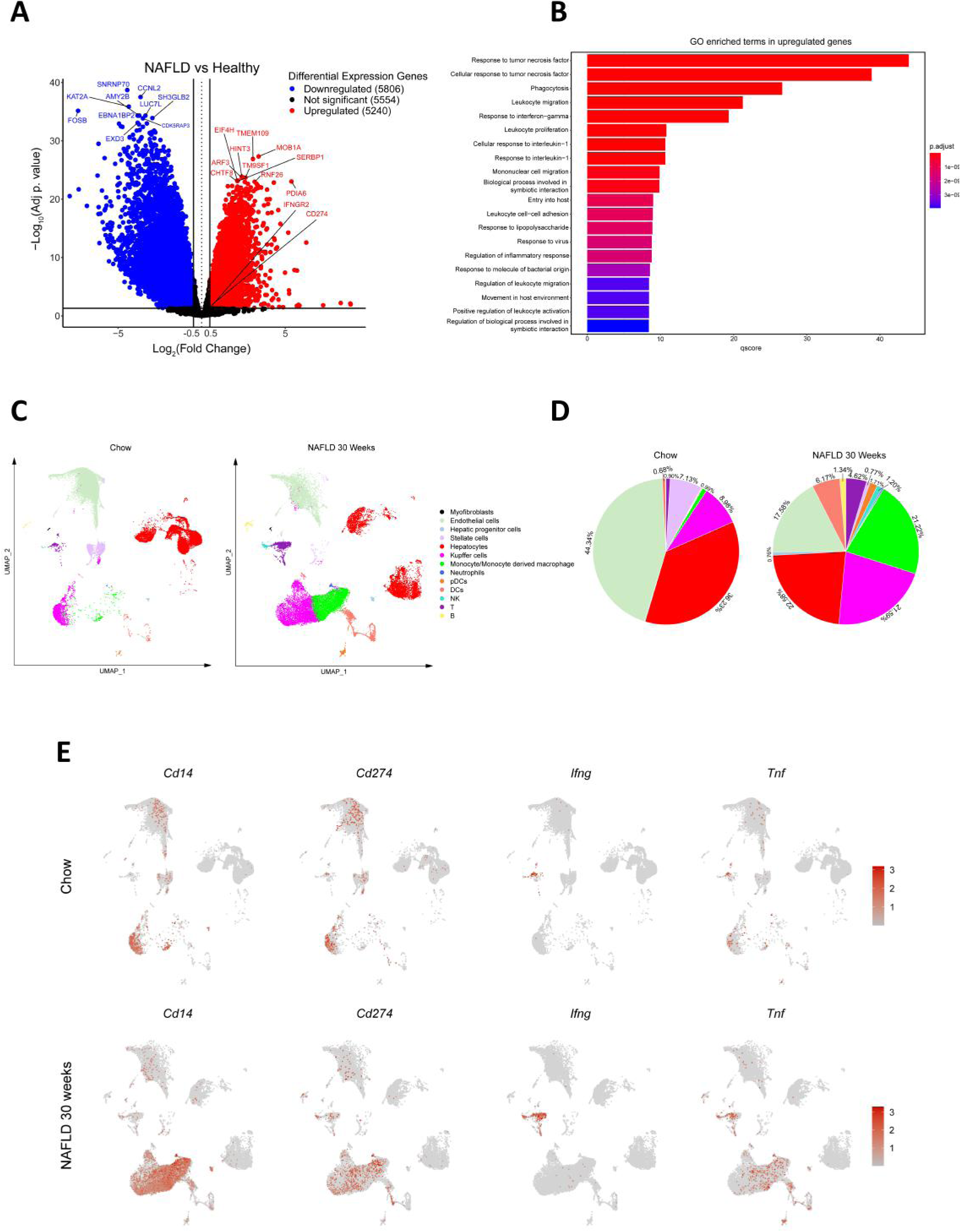
The expression of genes from the inflammatory pathways of IFN-γ and TNF-α is increased during the development of MAFLD in mice and humans. (A) Volcano plots illustrating the fold change and P-value for gene expression comparisons between livers of MAFLD and healthy patients. Genes of interest are highlighted on the volcano charts. (B) Gene Ontology (GO) functional analysis of differentially expressed genes (DEGs). GO enrichment analysis of DEGs was conducted using DAVID, presenting the 20 most significantly (P < 0.05) enriched GO terms in biological process, molecular function, and cellular component categories. All adjusted statistically significant values are presented as negative log-transformed base 10 values. (C) UMAP plots depicting immune cells from the livers of mice subjected to 30 weeks of HFCD diet compared to Chow diet. (D) Frequency distribution of immune cell populations. (E) Expression levels of CD14, CD274, IFN-γ, and TNF-α genes within hepatic immune cell populations.

Because IFN-γ and TNF-α signaling pathways are upregulated in MAFLD patients, we reanalyzed previously published single-cell RNA-seq data from liver samples from mice fed with a 60% high-fat diet, another known MAFLD model (22). We noticed an increase in immune cells in the hepatic tissue, such as monocytes, neutrophils, dendritic cells, T cells, and NK cells (Fig. 2C and 2D). Interestingly, we observed a specific upregulation of CD274 and TNF-α genes in monocytes and neutrophils, while T cells and NK cells showed increased IFN-γ expression (Fig. 2E), suggesting that the cytokines produced by these specific cells may be involved in the hyperinflammation observed during MAFLD.

Next, based on the results of increased IFN-γ and TNF-α signaling in bulk RNAseq and single-cell RNA sequencing analyses in hepatic samples of MAFLD patients and animals, we investigated the role of these cytokines in the susceptibility of endotoxemic MAFLD mice. Initially, we observed an exacerbated hepatic production of both cytokines induced by endotoxin challenge in the HFCD-fed group compared to the control mice treated with PBS or LPS (Fig. 3A and 3B). Furthermore, we also observed that IFN-γ^-/-^ and TNFR1R2α^-/-^ mice with MAFLD challenged with LPS did not show an increase in the enzyme ALT, a liver lesion marker (Fig. 3C and 3D). In addition, endotoxemic MAFLD TNFR1R2^-/-^ and IFN-γ^-/-^ mice exhibited reduced leukocyte infiltration in the liver tissue and improved survival (Fig. 3E, 3F, and 3G), respectively, as compared to endotoxemic WT mice. Furthermore, splenocytes from HFCD-fed IFN-deficient animals exhibited increased IL-6 and TNF-α production upon *in vitro* LPS stimulation, indicating a response pattern similar to that of control animals (Fig 2SA and Fig. 2SB). Together, our data demonstrate that MAFLD predisposes to the exacerbation of cytokine production, such as IFN-γ and TNF-α, which mediate an elevated index of mortality in endotoxemic mice.

**Figure 3.**
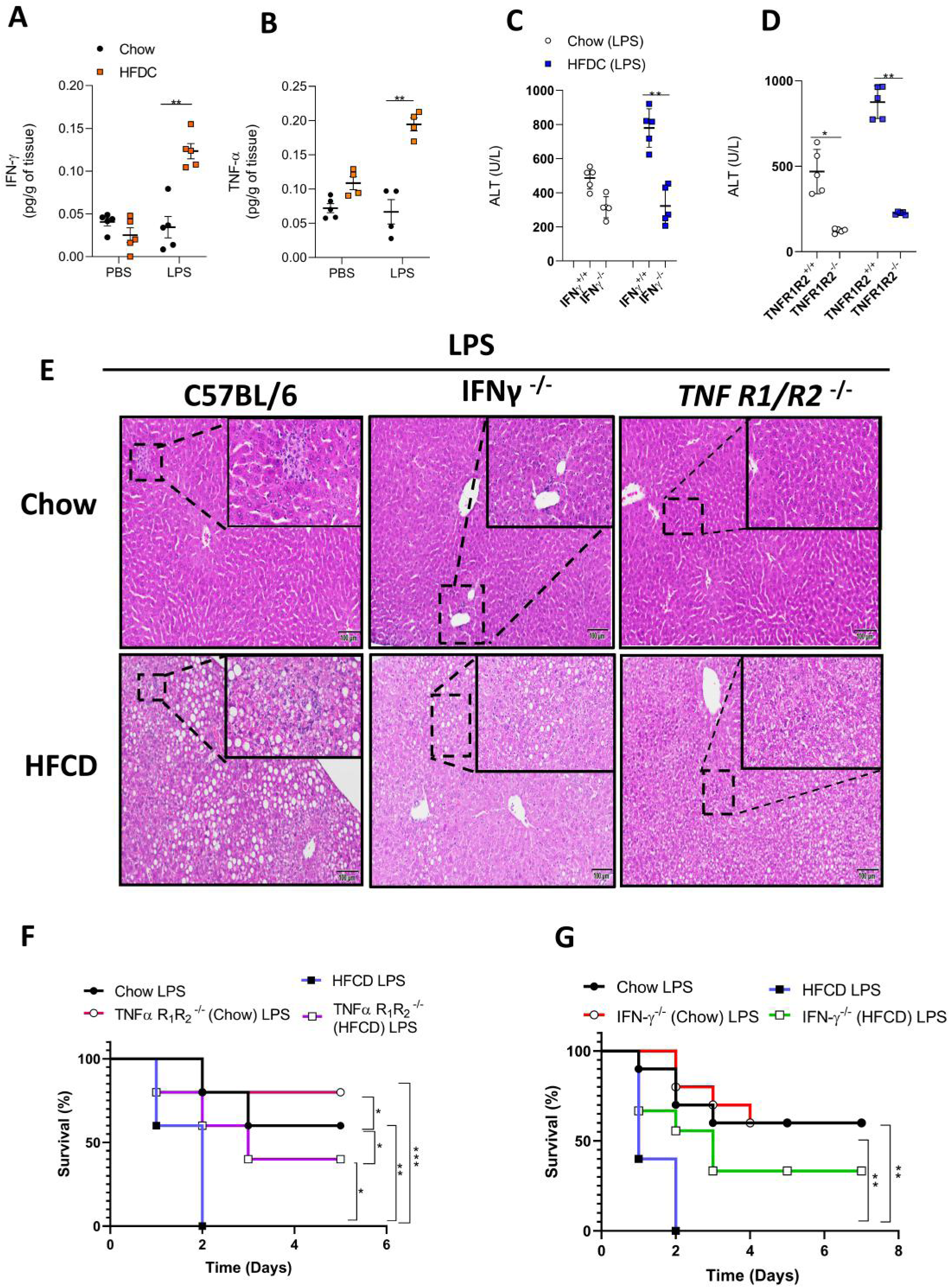
TNF-α and IFN-γ participate in susceptibility in mice with MAFLD to endotoxemia. (A) IFN-γ and (B) TNF-α secretion in the tissue livers of Chow and MAFLD mice injected with LPS (10mg/kg) (n = 5). (C) Serum ALT levels from mice with MAFLD and Chow and/or knockout of IFNγ (IFNγ-/-) and (D) TNF-α (TNFR1R2-/-), 6 hours after 10 mg/kg LPS and PBS administration (n=5). (E) Effect of LPS inoculation on liver histopathology from mice with MAFLD and Chow and/or knockout of IFNγ-/- and TNF-α (TNFR1R2-/-). (F) Survival curves of mice with MAFLD (n=5) and Chow (n=5) and/or knockout of TNF-α (TNFR1R2-/-) and (G) IFN-γ (IFN-γ-/-) after intraperitoneal inoculation of LPS (10mg/kg). The results are expressed as the mean ± standard deviation (SD) from one representative experiment. Statistical analysis was performed using one-way ANOVA followed by Tukey’s multiple comparison test. Statistical significance was determined with the following p-values: *p < 0.05, **p < 0.01, ***p < 0.001. Each experiment was independently repeated 2 times to ensure reproducibility of the findings.

### MAFLD induces the recruitment of NK cells that are sources of IFN-γ production in endotoxemia

During the inflammatory process, it is already established that CD4 and CD8 T cells are important sources of IFN-γ(23). Herein, we observed a selective increase in IFN-γ production by CD8, but not CD4, T cells during endotoxemia (Fig. 4A, Fig. 4B, Fig. S3A, Fig. S3B), respectively. In light of this, we next investigated the involvement of lymphoid cells in the high susceptibility of endotoxemic mice with MAFLD. To address this question, Rag1⁻/⁻ mice, which lack T and B cells, were subjected to the MAFLD protocol and then challenged with LPS, after which survival was assessed. Interestingly, *Rag1*-deficient animals under the HFCD remained susceptible to the LPS challenge (Fig. 4C) with exacerbation of liver injury (Fig. 4D), suggesting that these T and B cells do not contribute to susceptibility to the LPS challenge.

**Figure 4.**
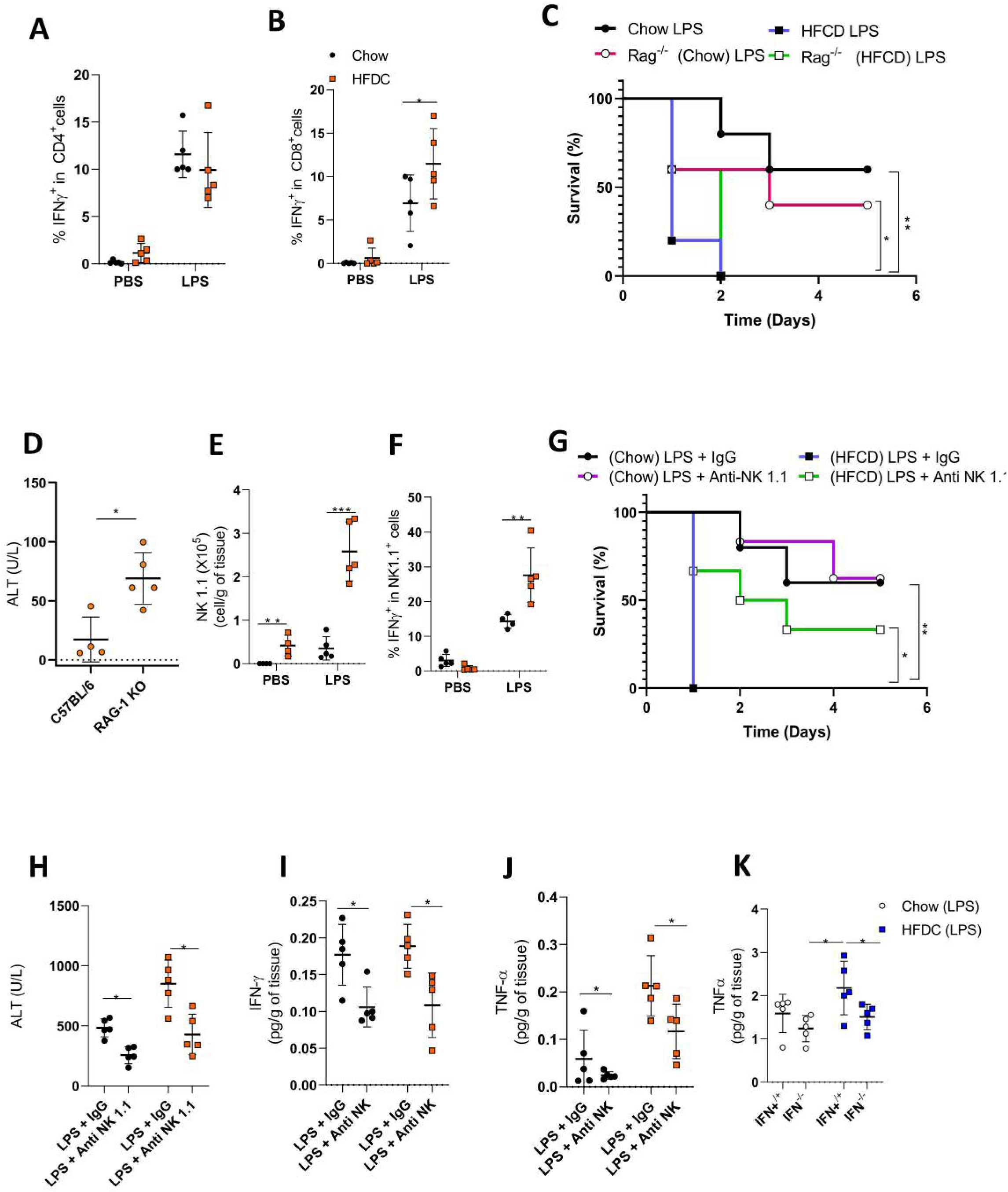
IFN-γ secreted by NK cells increases susceptibility of animals with MAFLD to endotoxemia. (A) CD4 IFN-γ^+^ and (B) CD8 IFN-γ^+^ T cell frequency in animals with MAFLD and Chow 6h after LPS inoculation. (C) displays survival curves of mice with MAFLD (n=5) and Chow (n=5) and T-cell knockout (RAg-/-). (D) Serum ALT levels from mice with MAFLD in knockout of Rag -/-, 6 hours after 10 mg/kg LPS (n=5). (E) and (F) represent the number of cells and frequency of NK IFNγ^+^ or IFN-γ^-^in animals with MAFLD and Chow 6h after LPS inoculation (G) The survival curves of mice with MAFLD (n=5) and Chow (n=5) and/or depleted NK cells (Anti NK 1.1), 6 days after intraperitoneal inoculation of LPS (10mg/kg) (H) presents the serum ALT levels from mice with MAFLD and Chow of mice with MAFLD (n=5) and Chow (n=5) and/or depleted NK cells (Anti NK 1.1), 6 hours after 10 mg/kg LPS or PBS administration (n=5). (I) and (J) depict IFN-γ and TNF-α secretion in the livers of Chow and MAFLD mice injected with LPS (10mg/kg) (n=5), respectively. (K) shows TNF-α secretion in the livers of Chow and MAFLD and/or knockout of IFN-γ (IFNγ-/-) mice injected with LPS (10mg/kg) (n=5).” The results are expressed as the mean ± standard deviation (SD) from one representative experiment. Statistical analysis was performed using one-way ANOVA followed by Tukey’s multiple comparison test. Statistical significance was determined with the following p-values: *p < 0.05, **p < 0.01, ***p < 0.001. Each experiment was independently repeated 2 times to ensure reproducibility of the findings.

Recent studies have demonstrated that NK cells are also IFN-γ–producing cells that are recruited to the hepatic tissue during MAFLD and that they are also involved in systemic inflammatory responses, including sepsis (24–29). We observed that NK cell numbers were increased in the liver of PBS-treated MAFLD mice when compared with mice fed a control diet (Fig. 4E). LPS challenge led to an increased accumulation of NK1.1⁺CD3⁻ NK cells in the liver of MAFLD mice, whereas NK1.1⁺CD3⁺ NKT cells were not detected (Fig. S3C and 4E). Furthermore, our cytometric analyses demonstrated that NK cells infiltrated in the liver tissue secrete high levels of IFN-γ (Fig.S3D and 4F). Next, to confirm the pathological involvement of NK cells during hyperinflammation in MAFLD, we depleted these cells by treating animals with anti-NK antibody 1.1, one day before the LPS challenge (Fig. S4). Interestingly, the pharmacological depletion of NK cells in MAFLD animals provoked an increased survival index (Fig. 4G). Furthermore, when NK cells were depleted, MAFLD animals exhibited reduced hepatic damage (Fig. 4H), IFN-γ levels (Fig. 4I), and TNF-α levels (Fig.4J). Together, these results suggest that NK cells are responsible for susceptibility via IFN-γ production in MAFLD during endotoxemia.

### Neutrophils are responsible for susceptibility during hyperinflammation in MAFLD

During MAFLD development, innate immune cells, such as monocytes and neutrophils, are the first leukocytes recruited to the liver (24, 30). With this in mind, mice with MAFLD were challenged with LPS, and the infiltration of these cells in the liver was quantified. During the initial inflammatory phase, six hours after the LPS challenge, we observed that monocytes were recruited to the site (Fig. 5A and 5B). Since inflammatory monocytes emigrate from the bone marrow towards the periphery via CCR2 and then migrate to inflamed tissues, differentiating into monocyte-derived macrophages (30, 31), we investigated the functional role of monocytes in MAFLD, using CCR2-deficient mice (Fig. 5S). Interestingly, animals deficient in monocyte migration (CCR2-/-) showed a high mortality rate compared to wild type after LPS challenge and neutrophil migration was not altered (Fig. 5SA and Fig. 5SB). Moreover, CCR2-/- animals subjected to the respective diets had leukocyte CD45+ infiltration similar to that observed in the LPS challenge control mice (Fig. 5SC and Fig. 5SD). Furthermore, the survival rate of MAFLD CCR2-/- mice challenged with LPS was not different of MAFLD wild type mice also challenged with LPS (Fig. 5SB). Together, the results suggest that monocytes do not contribute to susceptibility in endotoxemic animals with MAFLD

**Figure 5.**
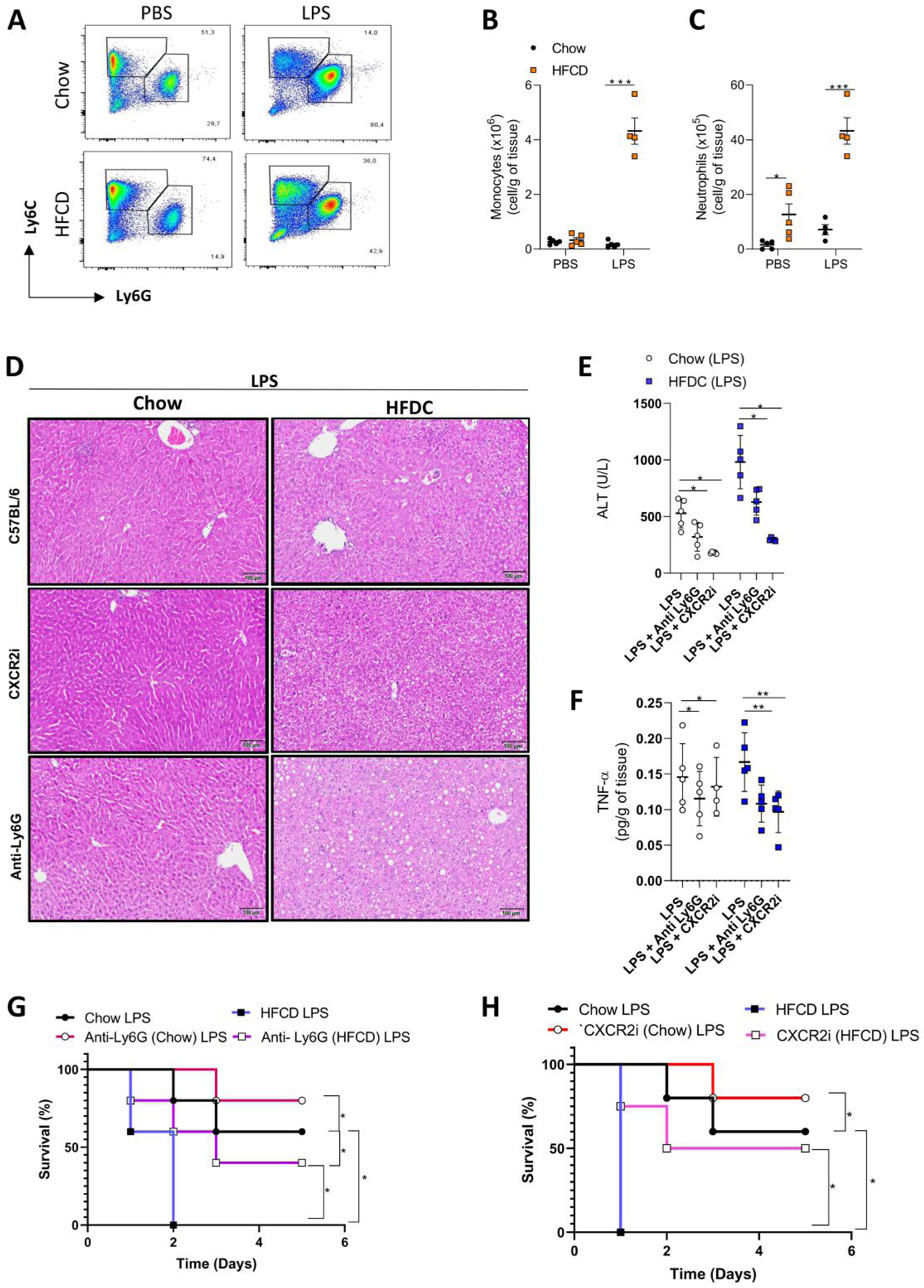
Neutrophils but not monocytes mediate susceptibility of animals with MAFLD to endotoxemia. (A) Representative flow cytometry plot showing monocytes (CD45+CD11b+MHC-Ly6C+Ly6G-) and neutrophils (CD45+CD11b+MHC-Ly6C+Ly6G+) and a number of monocytes (B) and neutrophils (C) in animals with MAFLD and Chow after LPS inoculation. (D) Effect of LPS inoculation on liver histopathology of mice with MAFLD and Chow treated with a CXCR2 inhibitor (CXCR2i) and neutrophil-depleted animals (anti-Ly6G). (E) Serum ALT levels of mice with MAFLD and Chow and/or CXCR2i and anti-Ly6G, 6 hours after administration of 10 mg/kg of LPS and PBS (n=5). (F) TNF-α secretion in the liver of Chow and MAFLD mice injected with LPS (10mg/kg) (n = 5). (G) Survival curves of mice with MAFLD (n=5) and Chow (n=5) treated with anti-Ly6G or (H) CXCR2i, and the survival rates were analysed daily for 6 days after intraperitoneal administration of LPS (10mg/kg). The results are expressed as the mean ± standard deviation (SD) from one representative experiment. Statistical analysis was performed using one-way ANOVA followed by Tukey’s multiple comparison test. Statistical significance was determined with the following p-values: *p < 0.05, **p < 0.01, ***p < 0.001. Each experiment was independently repeated 2 times to ensure reproducibility of the findings.

In addition to monocytes, granulocytes are innate immune cells recruited to the hepatic tissue during the development of MAFLD (19). We observed that high levels of neutrophils are present in the hepatic tissue of animals with MAFLD and that their recruitment is exacerbated in the hepatic microenvironment in endotoxemic mice (Fig. 5A and 5C). It is known that neutrophils are recruited to the inflammatory focus via CXCR2 (32). To understand the involvement of these receptors and neutrophils in MAFLD during endotoxemia, MAFLD animals were pre-treated one day before the LPS challenge with CXCR2i or with anti-Ly6G, which depletes these cells throughout the organism. Interestingly, after both treatments, we observed a reduction of the leukocyte infiltration into the hepatic tissue (Fig. 5D) and a decrease in hepatic lesions (ALT) (Fig. 5E) and TNF-α levels (Fig. 5F). Confirming the participation of granulocytes in susceptibility after the LPS challenge in MAFLD, animals treated with CXCR2i and anti-Ly6G showed an improvement in their survival (Fig. 5G and 5H).

### IFN-γ induces CD274 expression during endotoxemia in animals with MAFLD

IFN-γ is a cytokine that plays a crucial role in the T and NK cell-mediated immune response (27–29, 33–35). Recent studies have shown that this inflammatory cytokine also induces the expression of checkpoint molecules, such as programmed death-ligand 1 (PD-L1), in different immune cells, including neutrophils (30, 36, 37). Analyzing datasets from mice with MAFLD, we observed the upregulation of the CD274 (PD-L1) gene and TNF-α in neutrophils and IFN-γ in NK cells (Fig. 1C, Fig1D, and Fig. 1E). Considering these findings, we investigated whether IFN-γ induces the expression of PD-L1 in neutrophils in MAFLD mice during endotoxemia. First, we observed that only the MAFLD diet caused a significant increase in PD-L1 expression in CD45+CD11b+ leukocytes after LPS challenge (Fig. S6C). Analysis of this population revealed that neutrophils expressed higher levels of PD-L1 compared with monocytes (Fig. 6SA–6SD). Moreover, PD-L1⁺ neutrophils exhibited increased hepatic infiltration (Fig. 6A and 6B). Next, to confirm the role of IFN-γ in the expression of PD-L1, we investigated whether IFN-γ is capable of increasing the expression of PD-L1 in neutrophils after the LPS challenge of MAFLD. Mice with MAFLD lacking IFN-γ were challenged with LPS and assessed for PD-L1 expression in granulocytes. Interestingly, in the absence of IFN-γ, neutrophils do not express PD-L1, and their infiltration in the liver was attenuated (Fig. 6C and 6D).

**Figure 6.**
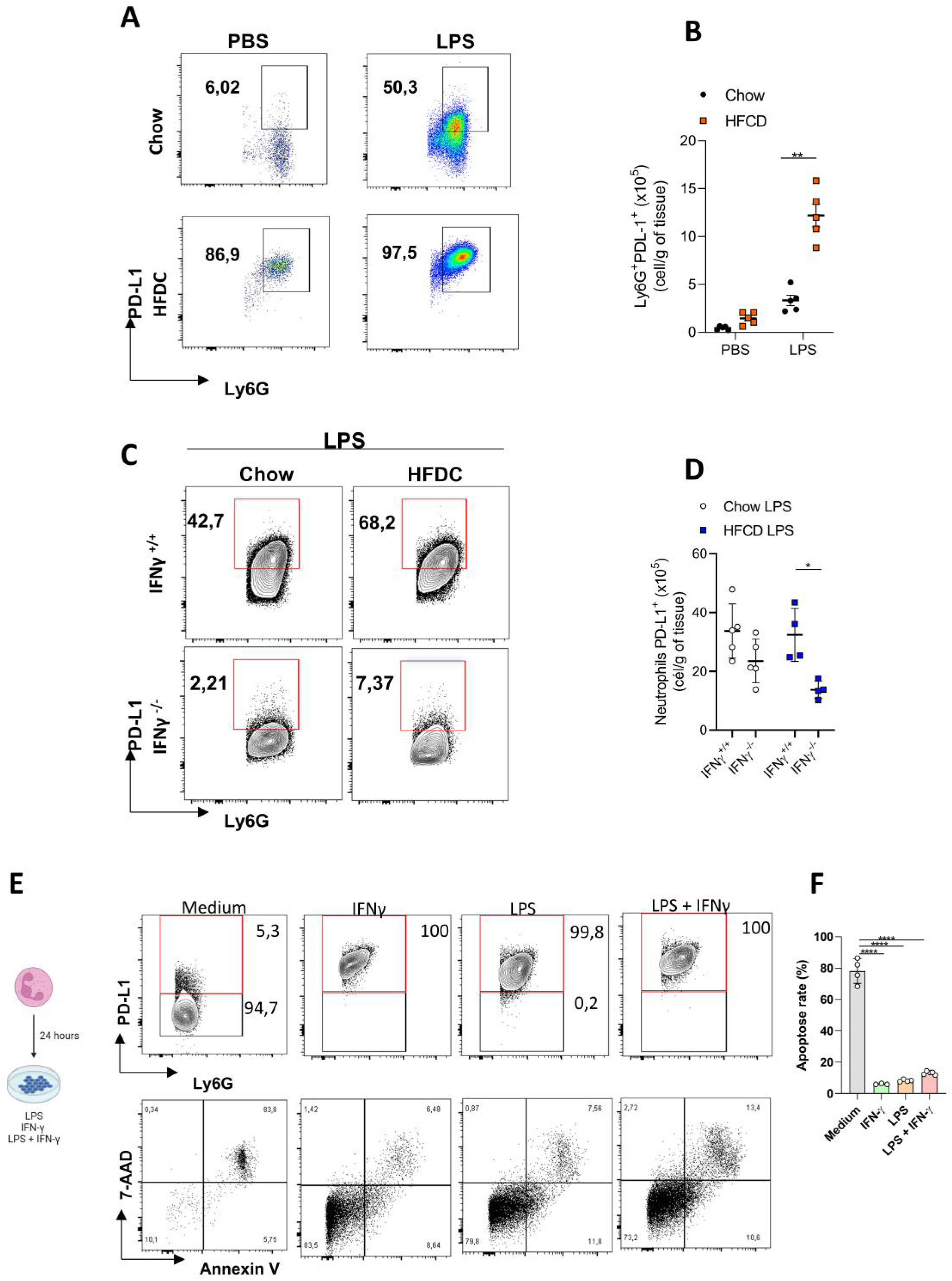
IFN-γ induces PD-L1 expression and reduces apoptosis in mouse neutrophils with MAFLD. (A) Representative flow cytometry plot showing the expression of PD-L1 in gated neutrophils and (B) the quantification of PD-L1+ neutrophils in the liver of mice with MAFLD or that received Chow diet, 6 hours after LPS inoculation (n=5 per group). (C) Representative flow cytometry plot showing the expression of PD-L1 in gated neutrophils and (D) the quantification of PD-L1+ neutrophils in the liver of mice with MAFLD and Chow and/or IFNγ knockout (IFNγ-/-) 6 hours after LPS inoculation (n=5 per group). (E) Representative flow cytometry plot showing the expression of PD-L1 and apoptosis (Annexin V+ 7AAD+) in gated neutrophils and (F) the frequency of apoptosis in neutrophils cultured in medium, LPS, IFN-γ, or LPS+ IFN-γ for 24 hours. The results are expressed as the mean ± standard deviation (SD) from one representative experiment. Statistical analysis was performed using one-way ANOVA followed by Tukey’s multiple comparison test. Statistical significance was determined with the following p-values: *p < 0.05, **p < 0.01, ***p < 0.001. Each experiment was independently repeated 2 times to ensure reproducibility of the findings.

As demonstrated previously in sepsis, PD-L1 neutrophils showed reduced rates of apoptosis (36). Therefore, we cultured neutrophils in the presence of IFN-γ and/or LPS and assessed their apoptosis rate. Interestingly, in these *in vitro* conditions, neutrophils expressing PD-L1 showed reduced rates of apoptosis (Fig. 6E and 6F), suggesting that PD-L1-expressing neutrophils can survive for long periods in the hepatic environment in MAFLD, accentuating the LPS-induced inflammatory response.

### Neutrophils expressing PD-L1+ secrete TNF-α during hyperinflammation in MAFLD

There is also evidence that excessive secretion of TNF-α by immune cells, especially neutrophils, may be associated with liver damage and the progression of MAFLD (36, 37). To investigate a possible link between the expression of PD-L1 by the neutrophils and their ability to express TNF-α, mice undergoing MAFLD were exposed to LPS, and the expression of TNF-α by PD-L1+ neutrophils was assessed. We observed an exacerbated expression of TNF-α by PD-L1+ neutrophils from MAFLD when compared to control animals (Fig. 7A, Fig. 7B, Fig. 7D and Fig8SG) and no increase in TNF-α^+^PDL1^-^ neutrophils (Fig. 7C). These results illustrate that after exposure of MAFLD animals to LPS, the expression of PD-L1 by neutrophils is linked to heightened TNF-α expression compared to control animals fed standard chow. To assess whether the lack of TNF-α signaling immunomodulates the granulocytic recruitment to the liver tissues in MAFLD animals, TNF receptor R1 and R2 knockout animals (TNF R1R2) were exposed to LPS. It was also observed that the absence of TNF-α did not interfere with the recruitment of neutrophils to the liver tissue (Fig. S8A and Fig. S8C) or the expression of PD-L1 in neutrophils (Fig. S8B and Fig. S8D). Similarly, the absence of TNF-α receptors in MAFLD during the challenge with LPS did not interfere with the production of IL-10 and IFN-γ (Fig. S8E and Fig. S8F). However, as described above, the absence of the TNF R1R2 receptor reduced liver injury and mortality induced by LPS administration in MAFLD (Fig. 3D and Fig. 3F).

**Figure 7.**
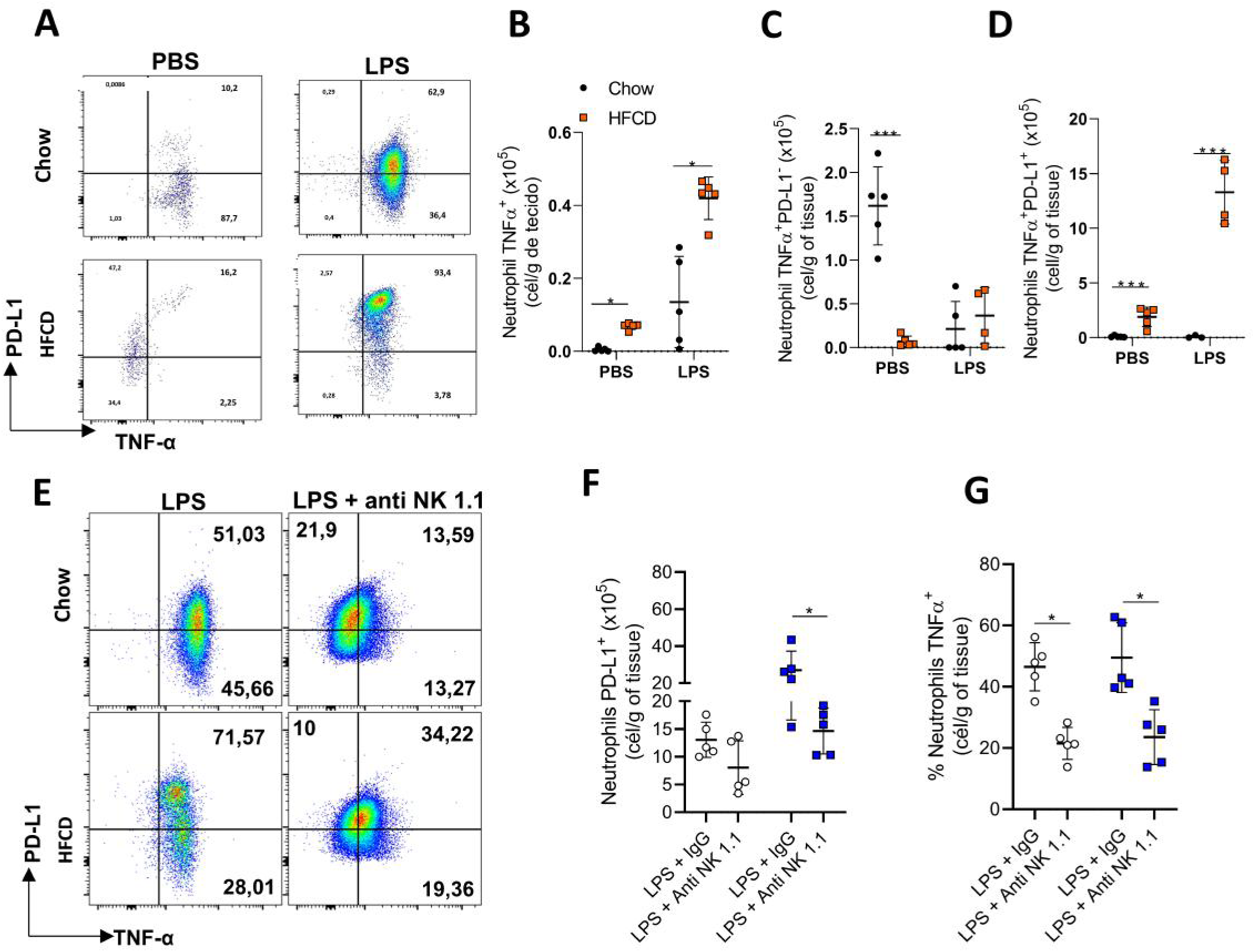
IFN-γ-dependent secretion of TNF-α by PD-L1+ neutrophils induces susceptibility of animals with MAFLD to endotoxemia. (A) Representative flow cytometry plot showing TNF-α expression in PD-L1+ neutrophils in the liver of animals with MAFLD or fed with Chow diet 6h after LPS inoculation. (B) Number of TNF-α+ neutrophils. (C) Number of TNF-α+PD-L1-neutrophils. (D) number of TNF-α+PD-L1+ neutrophils. (E) Representative flow cytometry plot showing TNF-α expression in PD-L1+TNF+ neutrophils in the liver of animals with MAFLD and Chow and/or depleted NK cells (anti-NK 1.1) 6h after LPS inoculation. (F) Number of PD-L1+ neutrophils. (G) Frequency of TNF-α+ neutrophils. The results are expressed as the mean ± standard deviation (SD) from one representative experiment. Statistical analysis was performed using one-way ANOVA followed by Tukey’s multiple comparison test. Statistical significance was determined with the following p-values: *p < 0.05, **p < 0.01, ***p < 0.001. Each experiment was independently repeated 2 times to ensure reproducibility of the findings.

Considering that NK cells present in liver tissue secrete IFN-γ (Fig. 4E) and that in IFN-γ^-/-^ animals, there is a reduction in TNF-α levels in the liver of mice with MAFLD during endotoxemia (Fig. 4K), we next examined whether PD-L1+TNF-α+ neutrophils are present in MAFLD animals depleted of NK cells. It was observed that in the absence of NK cells, PD-L1+TNF-α+ neutrophils were not detected (Fig. 7E, Fig. 7F, and Fig. 7G), confirming that IFN-γ secreted by NK cells is crucial for inducing PD-L1 expression and TNF-α secretion by neutrophils in the hepatic tissue in MAFLD animals after LPS administration.

### Anti-PD-L1 treatment protects animals with MAFLD during hyperinflammation

Next, we investigated the potential role of PD-L1 in TNF-α production by neutrophils and in the lesions in MAFLD mice. Initially, we found that anti-PD-L1, a biological drug inhibiting PD-L1’s interaction with PD-1, effectively prevented liver inflammation and tissue damage (Fig. 8A, Fig. 8B, and Fig. 8F). Additionally, this treatment reduced liver TNF-α levels without affecting IFN-γ (Fig. 8C) and improved host survival (Fig. 8D). Importantly, anti-PD-L1 did not decrease the number of recruited NK cells in all groups studied (Fig. 8I and Fig. 8L). Notably, treatment with anti-PD-L1 successfully blocked LPS-induced neutrophil recruitment and the increased presence of PD-L1-expressing neutrophils in mice on HFCD (Fig. 8J and Fig. 8K), while these parameters remained unchanged in mice fed a standard chow diet (Fig8G, Fig8H). Overall, our findings suggest that anti-PD-L1 therapy primarily suppresses exaggerated TNF-α production by neutrophils in MAFLD mice without affecting the IFN-γ secretory activity of NK cells mechanistically.

**Figure 8.**
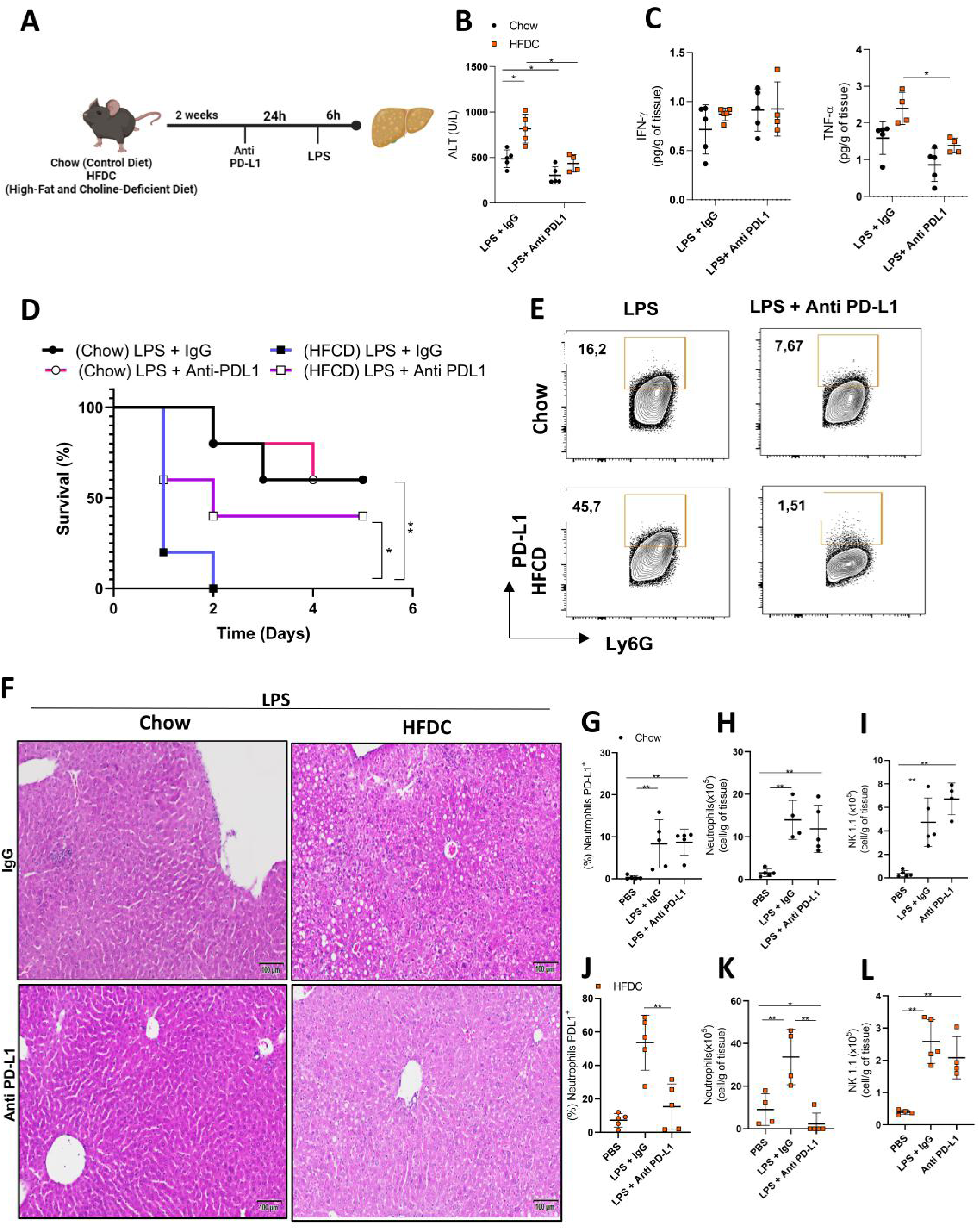
Anti PD-L1 treatment reduces PD-L1 neutrophils in animals with MAFLD during endotoxemia. (A) Schematic illustration of experimental design. (B) Serum ALT levels of mice with MAFLD or feded with Chow and/or treated with anti-PDL1, 6 hours after administration of 10 mg/kg of LPS or PBS (n=5). (C) IFN-γ and TNF-α secretion in the liver of Chow and MAFLD mice injected with LPS (10mg/kg) treated with anti-PDL1 (n = 5). (D) Survival curves of mice with MAFLD (n=5) and Chow (n=5) treated or not with anti-PD-L1. (E) Representative flow cytometry graph showing TNF-α expression in PD-L1+ neutrophils in the liver of animals with MAFLD and Chow and/or treated with anti-PD-L1 6h after LPS inoculation. (F) Effect of LPS inoculation in liver histopathology of mice with MAFLD and Chow treated or not with anti-PD-L1. (G and J) Frequency of PDL1 neutrophils, (H and K) Number of neutrophils, (I and L) Number of NK cells from animals with MAFLD and Chow and/or treated with anti-PDL1 after 6h of LPS inoculation. The results are expressed as the mean ± standard deviation (SD) from one representative experiment. Statistical analysis was performed using one-way ANOVA followed by Tukey’s multiple comparison test. Statistical significance was determined with the following p-values: *p < 0.05, **p < 0.01, ***p < 0.001. Each experiment was independently repeated 2 times to ensure reproducibility of the findings.

## 3. DISCUSSION

The pathogenesis of MAFLD and sepsis had been considered two independent conditions, associated with local and systemic inflammatory response, respectively. MAFLD is a chronic liver dysfunction characterized by the buildup of fat in the liver in individuals who do not consume excessive amounts of alcohol. On the other hand, sepsis is an inflammatory state triggered by an infection that leads to damage to vital organs, including the lungs and liver. This occurs when the body’s immune response fails to contain the microorganisms at the site of the primary infection(9). Recent studies demonstrate that the presence of MAFLD is associated with an increased risk of developing sepsis (38, 39), but the exact mechanisms underlying this association are not fully understood. In the present study, we observed that MAFLD mice induced by an HFDC diet are more susceptible to a systemic and hepatic hyper-inflammatory state induced by endotoxemia, confirming in our experimental condition that MAFLD can be considered a crucial risk factor of sepsis progression, even when MAFLD or hepatic inflammation is not yet detected. The LPS administration promotes the polarization of a specific subset of granulocytes expressing PD-L1, which secrete high levels of TNF-α in an NK-derived IFN-γ dependent manner. The released TNF-α plays an important role in the liver lesion. We also observed that CD4+ and CD8+ T cells or monocyte cells do not seem to participate in the mice’s susceptibility in this hyper-inflammatory state. In accordance, previous studies implicated NK cells in liver injuries in different MAFLD and sepsis experimental models(25, 40–44). It was observed that during severe sepsis, NK cell-released IFN-γ play a detrimental role in mediating tissue injury (44). Similarly, in MAFLD, NK cells can accumulate in the liver and release IFN-γ, which contributes to the development of hepatic inflammation and fibrosis(45). Therefore, these studies, support that the increase in NK cells induced by endotoxemia in MAFLD animals, along with IFN-γ secretion, appears to have a crucial role in the increased susceptibility of these pathological conditions, contributing to disease progression.

Considering this evidence, we re-analyzed previously published bioinformatic data (21, 22) and observed that, in MAFLD patients, IFN-γ and TNF-α are up-regulated, and that, in high-fat high fructose diet-fed mice, there is up-regulation of IFN-γ in NK cells, and TNF-α and CD274 (PD-L1) in myeloid cells (21, 22), suggesting that similar immune dysfunctions are observed in patients and experimental models of MAFLD. To add experimental information about how the IFN-γ mediates the liver lesion, we demonstrated that in endotoxemic MAFLD animals hepatic neutrophils are important producers of TNF-α, and this release is stimulated by NK-derived IFN-γ signaling. In addition, IFN-γ promoted the differentiation of PDL-1-expressing neutrophils, given that in those animals depleted of NK, the neutrophils did not exhibit this activated phenotype. Moreover, the PDL-1-expressing neutrophils showed a reduced apoptosis rate, increasing the retention time of the neutrophils in hepatic tissue. Thus, these events could explain the increase in hepatic injury in MAFLD animals during endotoxemia. According, in the context of infection-triggered inflammation, the participation of IFN-γ in modulating neutrophil pro-inflammatory profiles, including the expression of PD-L1, has been demonstrated (36). Neutrophils, as frontline innate immune cells, control pathogens through phagocytosis and the release of microbial mediators, including cytokines, cytosolic enzymes, free radicals, and NETs (46, 47). However, excessive neutrophil activity can lead to tissue damage, particularly in sepsis, where they contribute significantly to multi-organ damage via the overproduction of inflammatory mediators, including TNF-α (16, 48, 49). Our data demonstrate that during MAFLD, liver neutrophils are major sources of TNF-α production induced by endotoxemia. Confirming the involvement of TNF-α release by PDL-1+ neutrophils stimulated by NK cells-derived IFN-γ in the increase of susceptibility of MAFLD, we demonstrated that the absence of TNF-α signaling or depleted NK cells led to improved survival in mice with MAFLD during endotoxemia. However, the absence of TNF-α signaling did not affect PD-L1 expression in neutrophils. These findings imply that, while TNF-α plays a crucial tissue lesions, its intracellular signaling does not notably impair the expression of PD-L1 in the neutrophils or their migration into liver tissue during MAFLD. To reinforce that neutrophils are the main source of the TNF-α involved in susceptibility of MAFLD by endotoxemia, we evaluate whether the neutralization of these cells with specific antibodies could improve the survival of endotoxemic HFDC-fed mice. Numerous strategies for targeting neutrophils are being investigated in various clinical and experimental settings (12, 50), including the use of CXCR2 inhibitors in patients with NASH (13, 50). Interestingly, we demonstrated that targeting neutrophils with different biological drugs, similar to the inhibition of TNF-α, mitigated inflammation-induced liver lesions.

Recent studies suggest that during the progression of MAFLD, PD-L1 is expressed in both parenchymal and non-parenchymal cells (10, 51). Given that the neutrophils responsible for hepatic damage in endotoxemic MAFLD mice exhibit a specific PD-L1-expressing phenotype, *we* demonstrated that anti-PD-L1 significantly reduced TNF-α-producing PD-L1+ neutrophils in the liver environment, attenuating endotoxemia-induced inflammation and liver damage, ultimately reducing mortality in MAFLD mice. This highlights the role of PD-L1 expression in neutrophils in promoting a proinflammatory phenotype associated with heightened TNF-α production and susceptibility of endotoxemic HFDC-fed mice. In the context of endotoxemia, recent research indicates that PD-L1-expressing neutrophils also contribute to immunosuppression and increased susceptibility to secondary infections (52) and also contribute to lung injury. Thus, our data support that the inhibition of PD-L1, besides preventing the organ lesions induced by sepsis in MAFLD patients, could also prevent the long-term immunosuppression observed in these patients.

Although macrophages, including Kupffer cells, have been considered a relevant source of cytokines in endotoxemic animals (53, 54), we did not find an important role for these cells expressing CCR2 in the aggravation of endotoxemia, since the absence of these chemokine receptor, which governs the migration and differentiation of inflammatory monocytes into macrophages, did not influence the endotoxemia mortality in animals with MAFLD. However, it’s worth noting that we did not extensively investigate the interaction of IFN-γ and TNF-α in the role of monocytes in our model, necessitating further studies to unravel the possible mechanisms of these cells.

Our study presents several limitations that should be acknowledged and discussed. First, we cannot entirely rule out the possibility that our mice deficient in pro-inflammatory components exhibit reduced responsiveness to LPS. However, our ex vivo analyses using splenocytes from these animals revealed a preserved cytokine production following LPS stimulation. These results suggest that the in vivo differences observed are primarily driven by the MAFLD condition rather than by intrinsic defects in LPS sensitivity. Second, the absence of publicly available single-cell RNA-seq datasets from MAFLD subjects under endotoxemic or septic conditions limited our ability to perform direct translational comparisons. To overcome this, we analyzed existing MAFLD patients and experimental MAFLD datasets, which consistently demonstrated upregulation of IFN-y and TNF-α inflammatory pathways in MALFD. In line with these findings, our murine model revealed TNF-α⁺ myeloid and IFN-y⁺ NK cell populations, thereby reinforcing the validity and translational relevance of our results.

Overall, our findings establish a biological basis for understanding the progression of endotoxemia in MAFLD, highlighting the dysregulated role of neutrophils in which hepatic metabolic syndrome potentiates the inflammatory effects of granulocytes via NK cells, advocating interventions to mitigate this inflammatory condition.

## 4. MATERIAL AND METHODS

### Experimental Design

This study was designed to interrogate the pathologic mechanisms in MAFLD during endotoxemia-induced generalized inflammation. The mice were submitted to the HFDC diet for two weeks, and subsequently, LPS administration and the cellular mechanisms were outlined after 6h of the challenge, knowing that no animal succumbed to death (indicated by survival). Furthermore, knockout animals and those treated with pharmacological inhibitors or neutralizing antibodies shared the same control groups (chow and HFCD), as required by the animal ethics committee. Inflammatory infiltrates along with liver lesions were assessed based on H&E-stained tissue sections, and endpoint analyses included flow cytometry.

### Animals

The mice used in this study were in the C57BL/6 genetic background and included CCR2 ^-/-^, IFN-γ^-/-^, and TNFR1R2 ^-/-^. Male mice aged 5 to 6 weeks were reared and maintained under specific pathogen-free conditions in the University of São Paulo vivarium, FMRP/USP. All animals received food and water ad libitum, at 25°C, with food and water ad libitum. Mice were fed a standard low-fat diet (hereinafter referred to as the standard “Chow” diet – 10% of calories derived from fat) or a Choline Deficient High Fat Diet (referred to as “HFCD”, 60% of calories derived from fat and choline-deficient) for 2 weeks for the development of MAFLD. The diets were provided by Rhoster (Araçoiaba da Serra- São Paulo). Care for the mice complied with institutional guidelines on ethics in animal experiments; approved by CETEA (Ethics Committee for Animal Experimentation of the Faculty of Medicine of Ribeirão Preto, approved protocol number 193/2019).

### Endotoxemia

Lipopolysaccharide (LPS; Escherichia coli (O111:B4), L2630, Sigma-Aldrich, St. Louis, MO, USA) was administered intraperitoneally (i.p.; 10 mg/kg) in C57BL/6, CCR2 -/-, IFN-γ-/-, and TNFR1R2 -/- mice. The HFCD was initiated in 5–6 week-old mice, and LPS was administered after 2 weeks on the diet, meaning that LPS administration occurred when the mice were 7–8 weeks old, with body weights ranging from 22 to 26 g. LPS was previously solubilized in sterile saline and frozen at -70°C. The animals were euthanized 6 hours after LPS administration.

### Survival analysis

Survival was monitored daily, and survival curves were generated using the Kaplan–Meier method. Experimental groups were composed of five animals per group (n = 5), in accordance with the requirements of the institutional animal ethics committee and the use of shared control groups. Sample size was defined based on an a priori power analysis, which indicated that this number of animals per group was sufficient to detect biologically relevant differences in survival with adequate statistical power. Survival distributions among groups were compared using the log-rank (Mantel–Cox) test. Statistical significance was considered when *p* < 0.05.

### Hepatic non-parenchymal cell isolation

Livers were collected, and 200-300 mg of tissue was taken for flow cytometry analysis. The tissue was minced with surgical scissors and then digested with 1 mg of collagenase II, and 60 U/ml of DNase I in PBS at 37 °C for 45 minutes, shaking at 110 rpm. The digested tissue was then processed through a 100 µm cell strainer and centrifuged at 1,500 rpm for 5 minutes. Subsequently, the isolation of non-parenchymal cells was performed by Percoll (GE Healthcare) centrifuged at 2300rpm at 22 °C for 30 minutes, subjected to ACK (red blood cell lysis), and resuspended in 0.5% BSA in PBS for staining. Single-cell suspensions were stained with Zombie Aqua viability stain (Biolegend), blocked with TruStain FcX (Biolegend), and then stained with flow cytometry antibodies when indicated; antibodies are tabulated in Table S1. As animals were already in a hyperinflammatory state, no additional in vitro stimulation was required. Intracellular detection of IFN-γ and TNF-α was conducted using the Foxp3 staining kit (eBioscience). Since both antibodies were conjugated to PE, staining and analyses were performed in separate experiments. Cells were collected on the CANTO II flow cytometer using BD FACS Diva software and then analyzed using FlowJo (USA).

### Histology

Mouse livers were fixed in 10% formalin overnight, washed twice with PBS, and placed in 70% ethanol or PBS until processing. For H&E, tissues were embedded in paraffin, sectioned, and stained with H&E at the MGH Histopathology Research Core.

### Cytokine quantification

For the detection of IFN-γ, TNF-α, and IL-10 in the liver, tissue samples were weighed and titrated in 1 mL of PBS Complete (Roche Diagnostics, Mannheim, Germany) containing a cocktail of protease inhibitors. Cytokine levels were determined using commercial ELISA kits for IFN-γ, TNF-α, and IL-10 (Biolegend, USA). Wavelength correction and background signals were subtracted from absorbance values

### Serum analysis

At the time of sacrifice, blood samples were collected in EDTA collection tubes and subsequently centrifuged to obtain serum. Serum ALT and Urea levels were measured using a colorimetric kit (BioQuant, Heidelberg, Germany) according to the manufacturer’s instructions.

### Neutrophil targeting

For neutrophil targeting, antibodies/inhibitors were used as follows: Anti-Ly6G (BioXCell, catalog # BP0075) was administered at 500 mg/kg per mouse on day 13 after starting the diet (24h before inoculation of the LPS) and sacrificed on day 14. For survival studies, all mAbs were inoculated once daily for 7 days. CXCR2 inhibitor SB 225002 (Tocris, catalog #2725/10) was injected at 200 mg per dose and programmed as anti-Ly6G. For anti–PD–L1–based neutrophil targeting, anti–PD-L1 (BioXCell, catalog #BE0101) was injected at 12.5 mg/kg ∼12h before the LPS challenge and again 1 day after treatment, each time followed by anti-mouse IgG2b (12.5 mg/kg) (BioXCell, catalog no. BE0252). All injections were intraperitoneal.

### Neutrophil purification and apoptosis assay

Peripheral blood samples were collected by venipuncture and centrifuged at 450 × g for plasma separation. The blood cells were then resuspended in Hank’s balanced salt solution (Corning; cat. 21-022-CV), and the neutrophil population was isolated by Percoll density gradient (GE Healthcare; cat. 17-5445-01) (72 %, 63%, 54% and 45%). Isolated neutrophils were resuspended in RPMI1640 (Corning; cat. 15-040-CVR) supplemented with 0.5% BSA. Neutrophil purity > 95% was determined by Rosenfeld color Cytospin (Laborclin; cat 620529)

To quantify apoptosis, cells were permeabilized with Triton X-100 and stained with propidium iodide. After 10 minutes of incubation at 4 degrees in the dark, neutrophil apoptosis was quantified as PI+ staining on FACSCanto II (Becton Dickinson, Franklin Lakes, NJ, USA). Results were analyzed using Flowjo software (Tree Star, Ashland, OR, USA). Apoptosis was detected as a measure of the hypodiploid peak in PI fluorescence histograms, as described by Nicoletti et al. BD Bioscience, Franklin Lakes, NJ, USA) and Annexin V and 7-AAD double staining (Biolegend, San Diego, USA).

### Splenocyte culture

Mice were fed a control chow or HFCD diet for 2 weeks. Animals were euthanized, and spleens were aseptically removed in cold PBS (Gibco) and then mechanically dissociated through a 100-µm cell strainer to obtain a single-cell suspension. Red blood cells are lysed with ACK buffer for 1–2 min at room temperature, the reaction is quenched with complete RPMI 1640 medium (supplemented with 10% heat-inactivated FBS, 2 mM L-glutamine, and antibiotics), and cells are centrifuged and resuspended at the desired density, 1 × 10^6^ cells/mL. Splenocytes are plated in 96-well plates and incubated at 37°C, 5% CO₂. To induce immune activation, cells are stimulated with lipopolysaccharide (E. coli O111:B4) at concentrations ranging from 100 ng/mL to 1 µg/mL for 24 h, and cytokine dosage was performed for IL-6 and TNF-α.

### NK cell depletion

For in vivo NK cell depletion, male C57BL/6 mice received NK cell neutralizing antibody (BioXCell, catalog # PK136) or isotype-matched mouse IgG2α monoclonal antibody control (BioXCell, catalog # BE0252) by intraperitoneal injection (150 μg in 200 μL PBS 24h before LPS challenge). To assess survival, the antibodies were inoculated 24 hours before the LPS challenge and then once a day for 7 days.

### Single-cell RNA sequencing analysis

We re-analyzed single-cell transcriptomic data (GSE166504) from the mouse liver from mice with MAFLD and their respective control groups (21). The dataset was downloaded, and the RDS file was imported into R (55) environment version v4.2.3 and Seurat v4.1.1 (56) by filtering genes expressed in at least 300 cells. For the pre-processing step, outlier cells were filtered out based on three metrics (library size < 20,000, number of expressed genes between 200 and 4,000, and mitochondrial percentage expression < 0,8). The top 3,000 variable genes were then identified using the ‘best’ method using the FindVariableFeatures function. Percent of mitochondrial genes was regressed out in the scaling step, and Principal Component Analysis (PCA) was performed using the top 3,000 variable genes, and the top 30 PCs were selected for dimension reduction by Uniform Manifold Approximation and Projection (UMAP). Clusters were identified using the author’s annotation. Then, differential gene expression analysis was performed using the FindAllMarkers function in Seurat with default parameters to obtain a list of significant gene markers for each cluster of cells. Visualization of genes illustrating expression levels was performed using R/Seurat commands (DimPlot, FeaturePlot, and DotPlot) using ggplot2 (57) and customized (58) R packages.

### RNA sequencing analysis

RNA-seq data were obtained from GSE185051 (59). The R/Bioconductor package edgeR was used to identify differentially expressed genes among the samples, after removing absent features (zero counts in more than 75% of samples) (60). Genes with a fold change of >0.5 were identified as differentially expressed.

### Pathway enrichment analysis

The list of differentially expressed genes was enriched using the ClusterProfiler R package (8). Gene ontology (GO) terms in the Biological Processes category with P < 0.05 were considered significant. Statistically significant, non-redundant GO-enriched terms were plotted.

### Statistics

Statistical significance was determined by two-tailed paired or unpaired Student’s t-test or one-way ANOVA followed by Tukey’s post hoc test. Absolute numbers and percentages were compared using Fisher’s exact test. Spearman rank order correlation (r) was calculated to describe the correlations. P<0.05 was considered statistically significant. Statistical analyses and graphics were performed using the GraphPad Prism 8.4.2 software.

## Author contributions

CCOB, AK, and FQC designed the study. CCOB, AK, PSAA and GCMC performed the mouse experiments. CCOB, AK, and FQC wrote the manuscript. GVLS performed the bioinformatic analysis. CCOB, AK, LOL, TMC, JCFAF, and FQC reviewed the manuscript.

## Acknowledgements

We thank members of Group FQC for their advice and clinical assistance, and Marcella, Sergio R. Rosa, Katia Santos, Ieda Schivo for technical assistance.

