## Supplementary Figure 1 for "PDL-1+ Neutrophils mediate susceptibility during endotoxemia in Metabolic Dysfunction–Associated Steatotic Liver Disease"

**Figure S1. Validation of the HFCD-induced MAFLD experimental model based on liver weight, blood glucose levels, and histological analysis (H&E and Picro Sirius staining).**

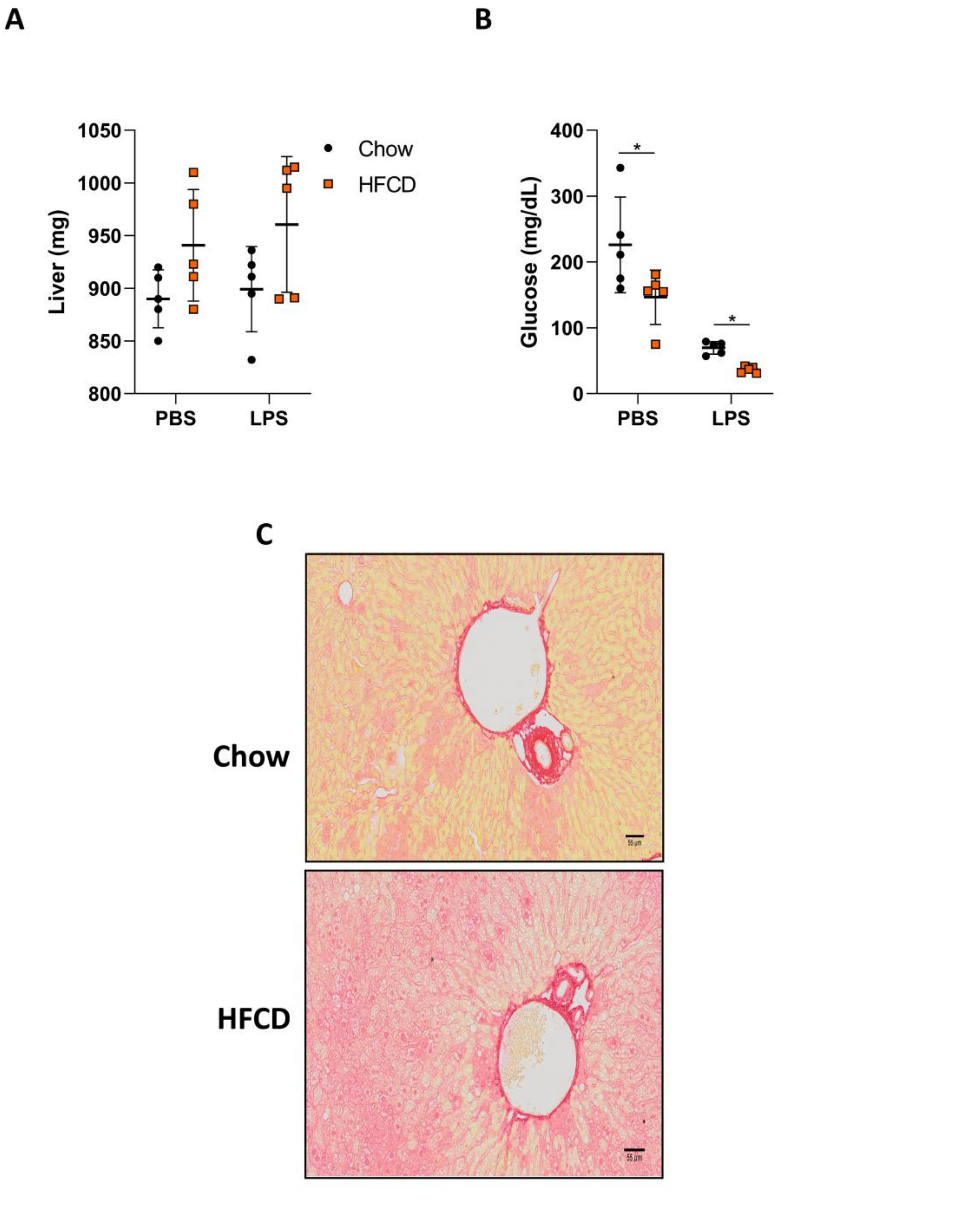
