## Supplementary figures and images for "PDL-1+ Neutrophils mediate susceptibility during endotoxemia in Metabolic Dysfunction–Associated Steatotic Liver Disease"

### Supplementary Figure 2

**Figure S2. The IFNy-deficient mice display LPS responses comparable to controls.**

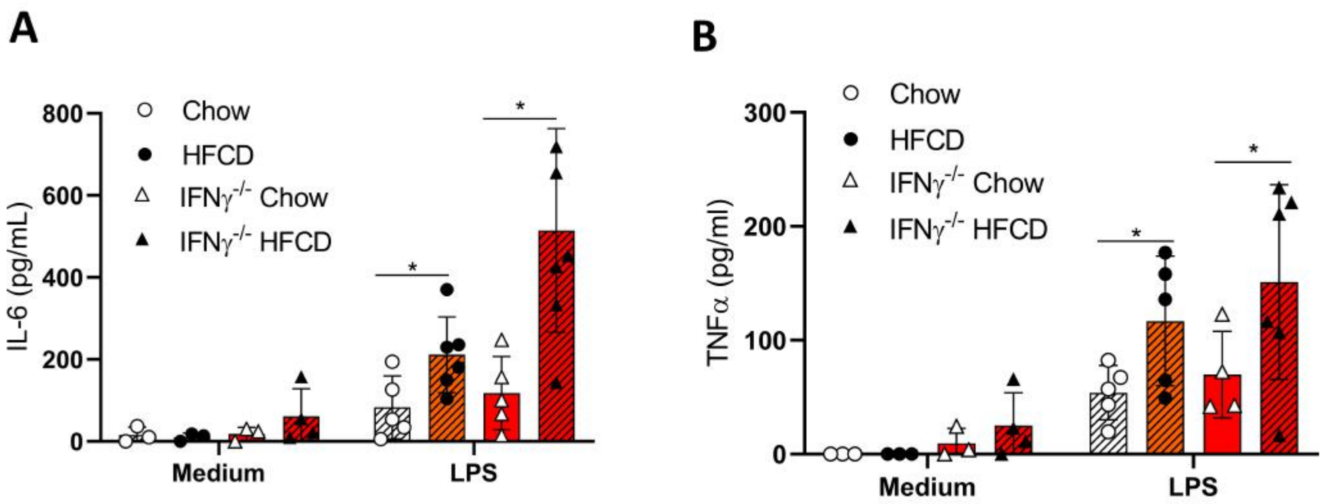

### Supplementary Figure 3

**Figure S3. Nk cells produce high levels of IFN during endotoxemia in MAFLD**

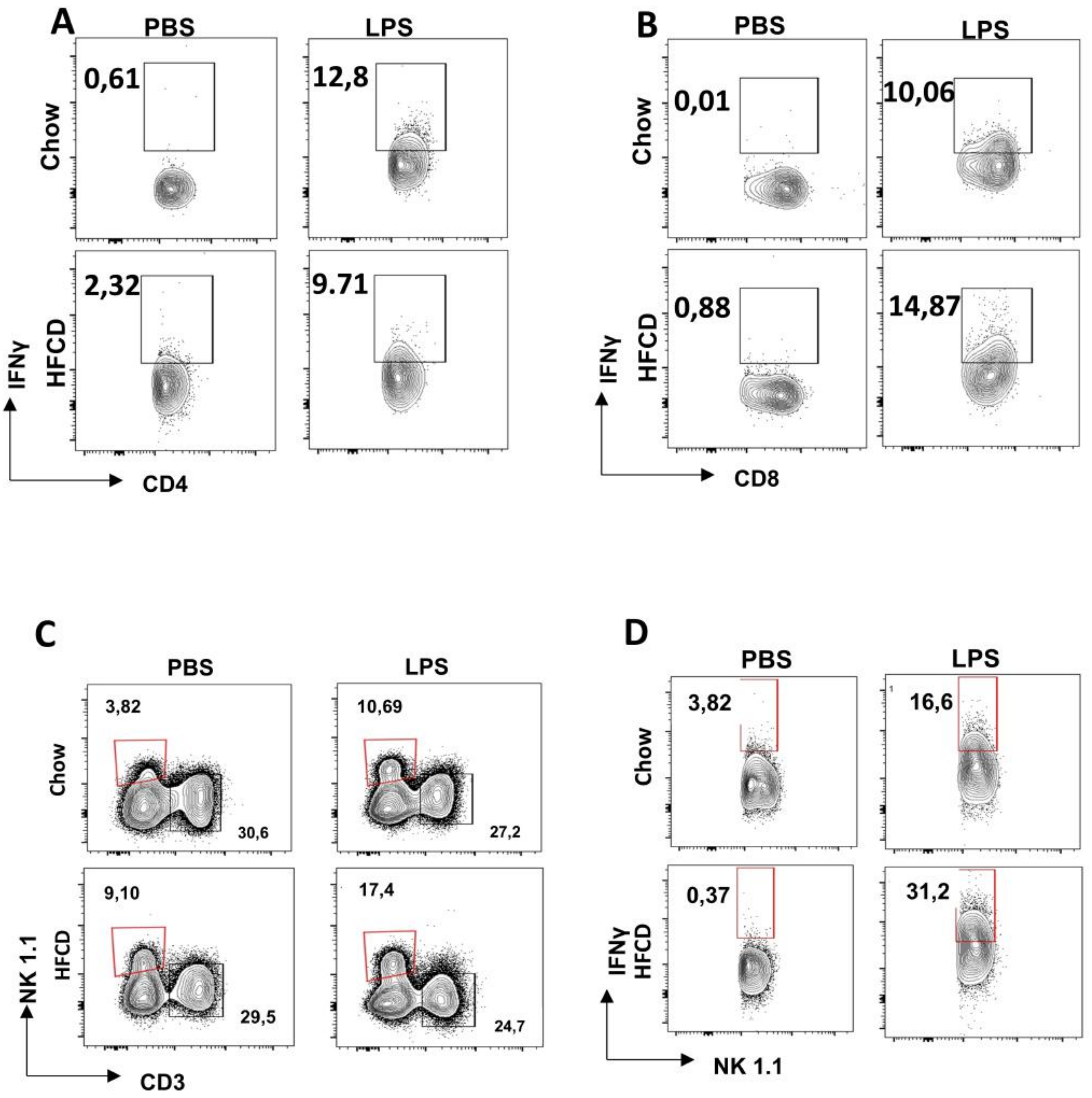

### Supplementary Figure 4

**Figure S4. Anti-NK 1.1 treatment deplete NK cells in animals with MAFLD during endotoxemia**

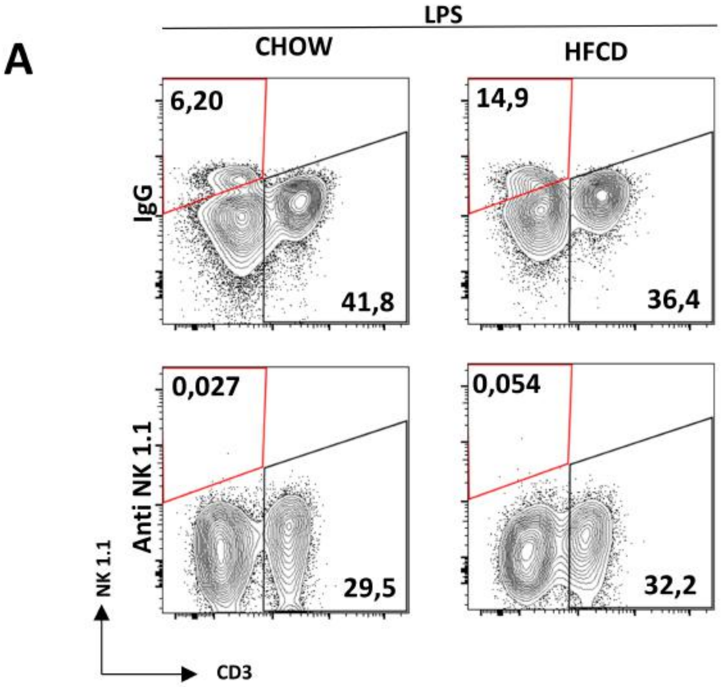

### Supplementary Figure 5

**Figure S5. Monocytes do not participate in susceptibility in mice with MAFLD to endotoxemia**

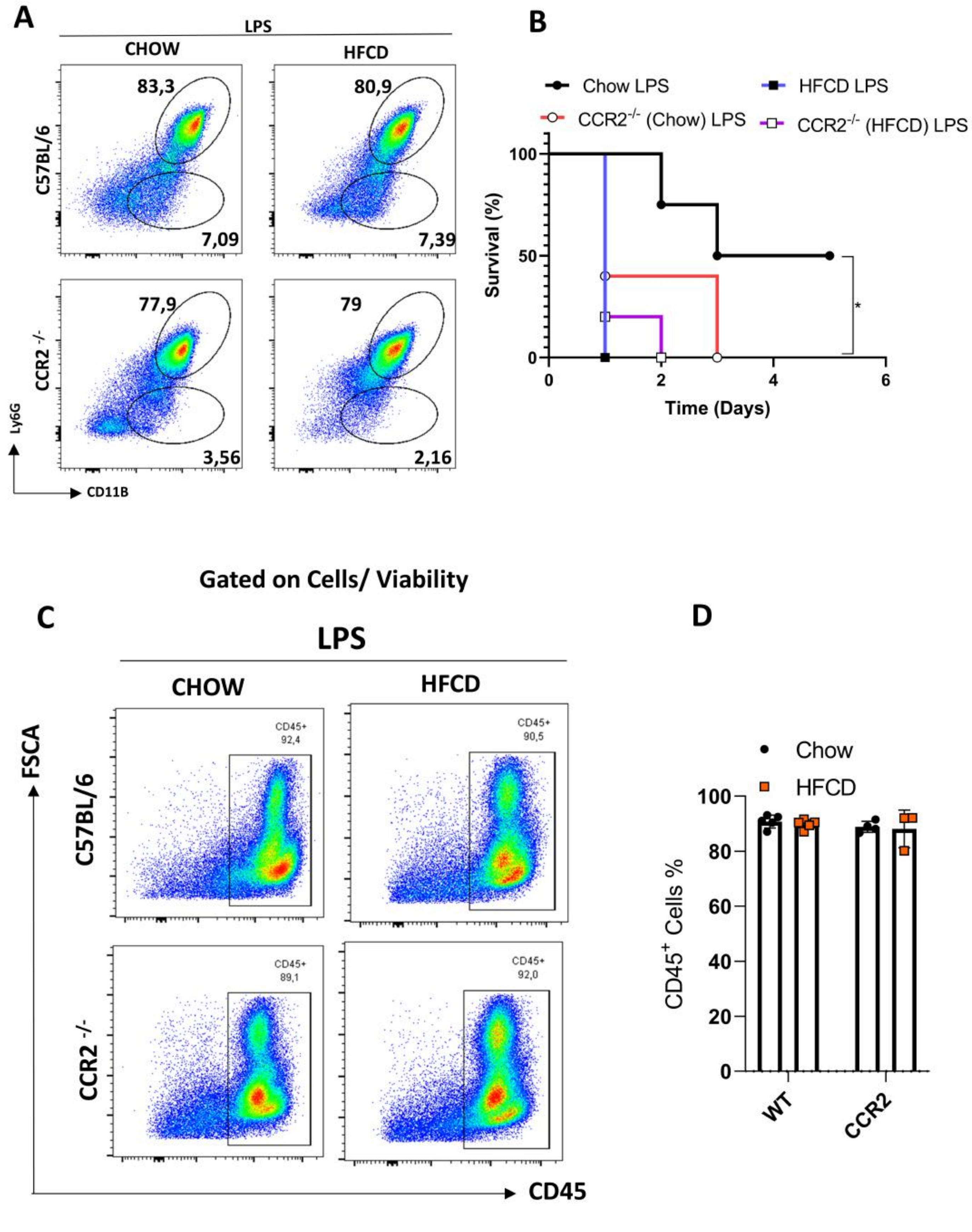

### Supplementary Figure 6

**Figure S6. Neutrophils are the main PD-L1–expressing subset among $CD45^{+}CD11b^{+}$ leukocytes**

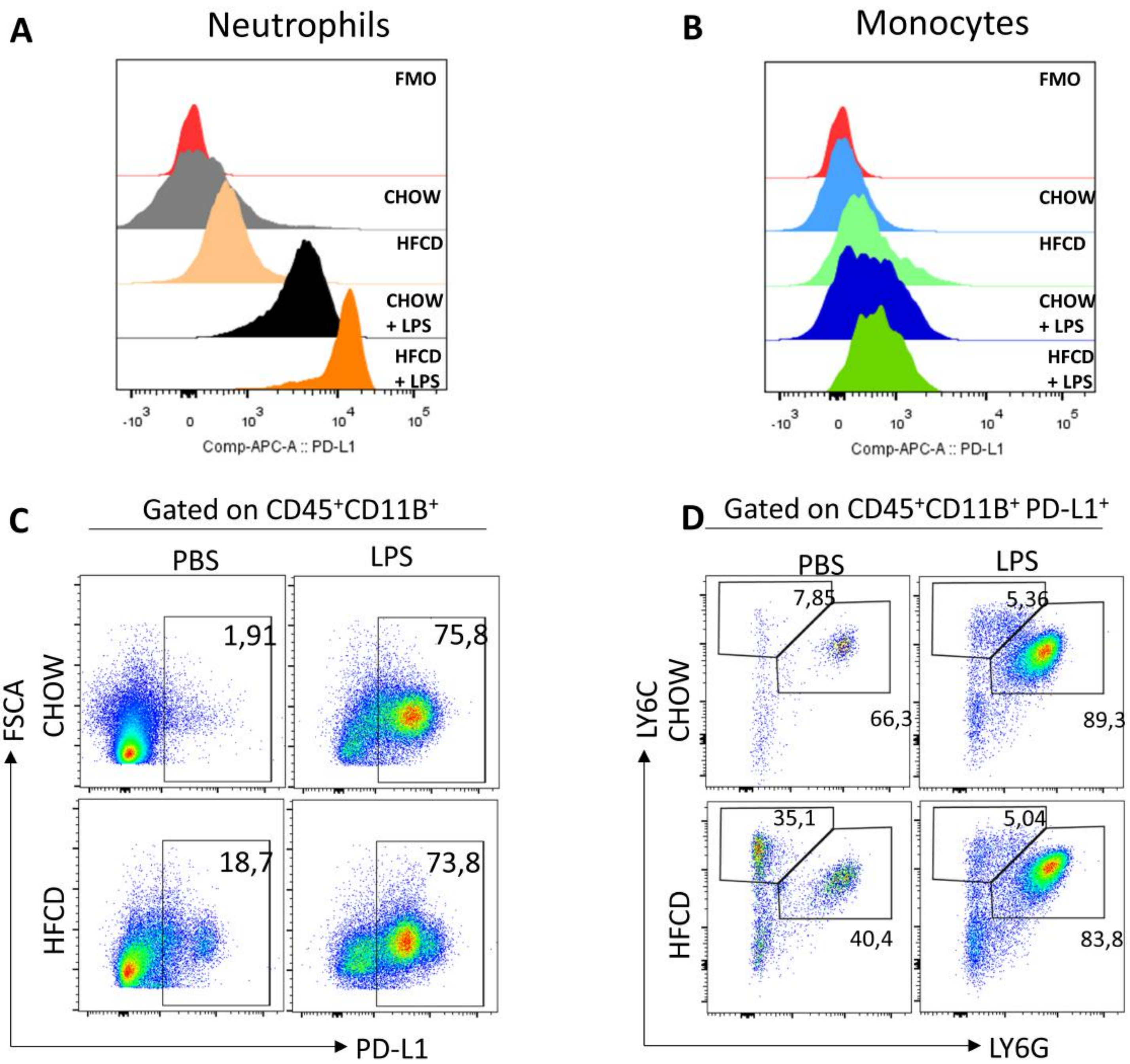

### Supplementary Figure 7

**Figure S7. CXCR2i treatments reduce IFN-γ in liver tissue in animals with MAFLD to endotoxemia**

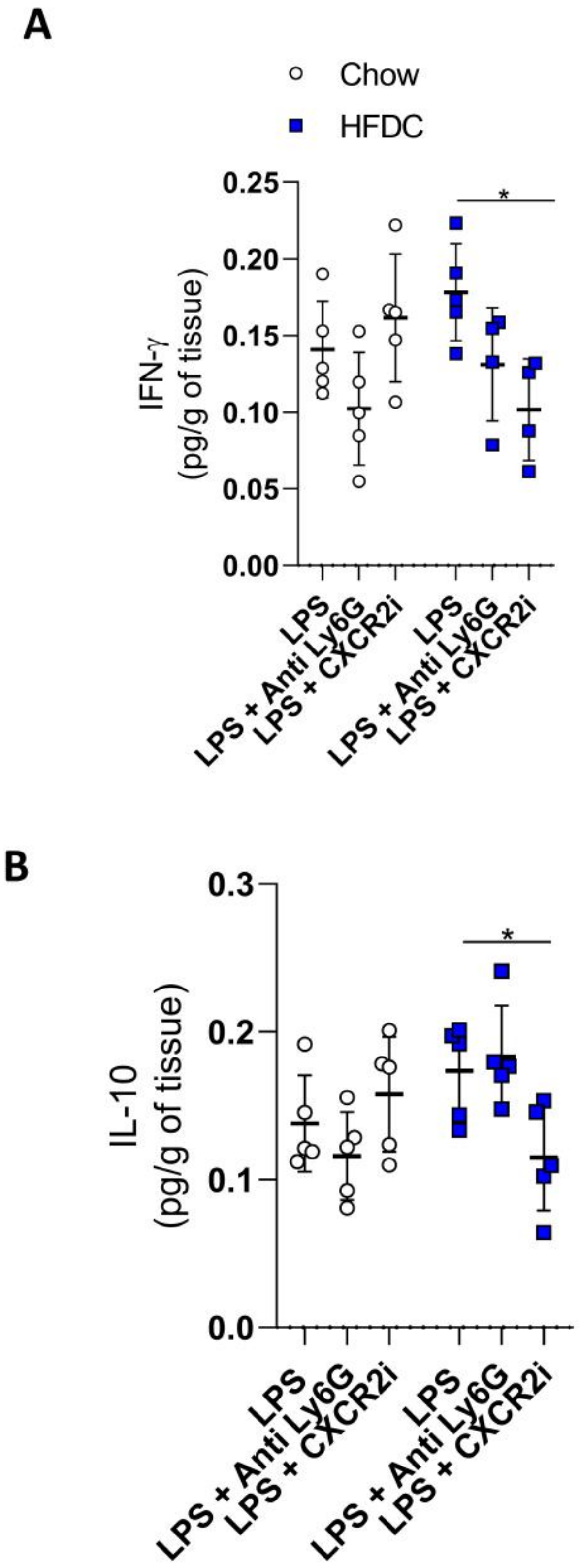

### Supplementary Figure 10

**Figure S10. Gating strategy for identification of NK, CD4 and CD8 T cells**

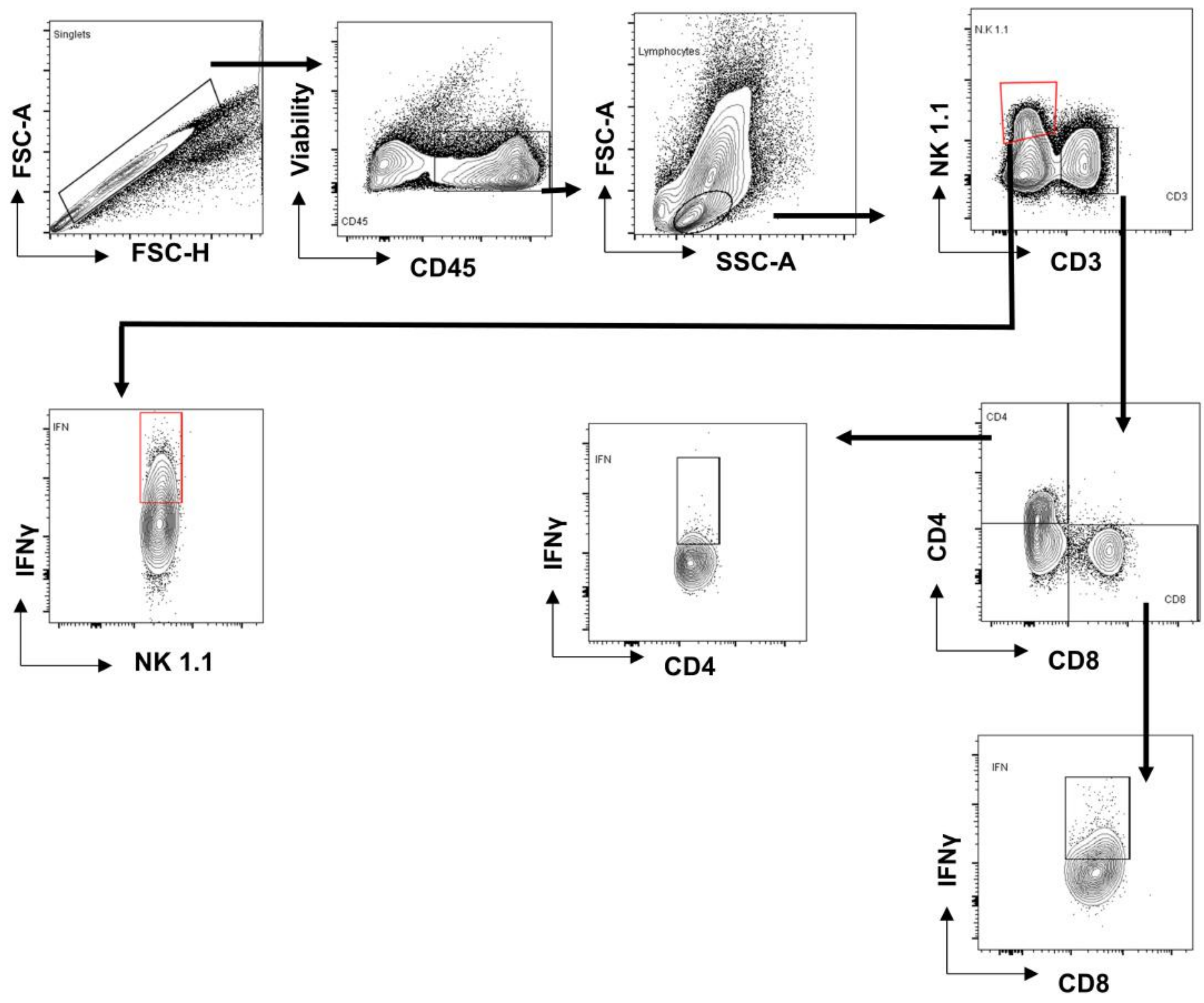

### Supplementary Figure 11

#### Figure S11. Gating strategy for identification of PD-L1+ neutrophils

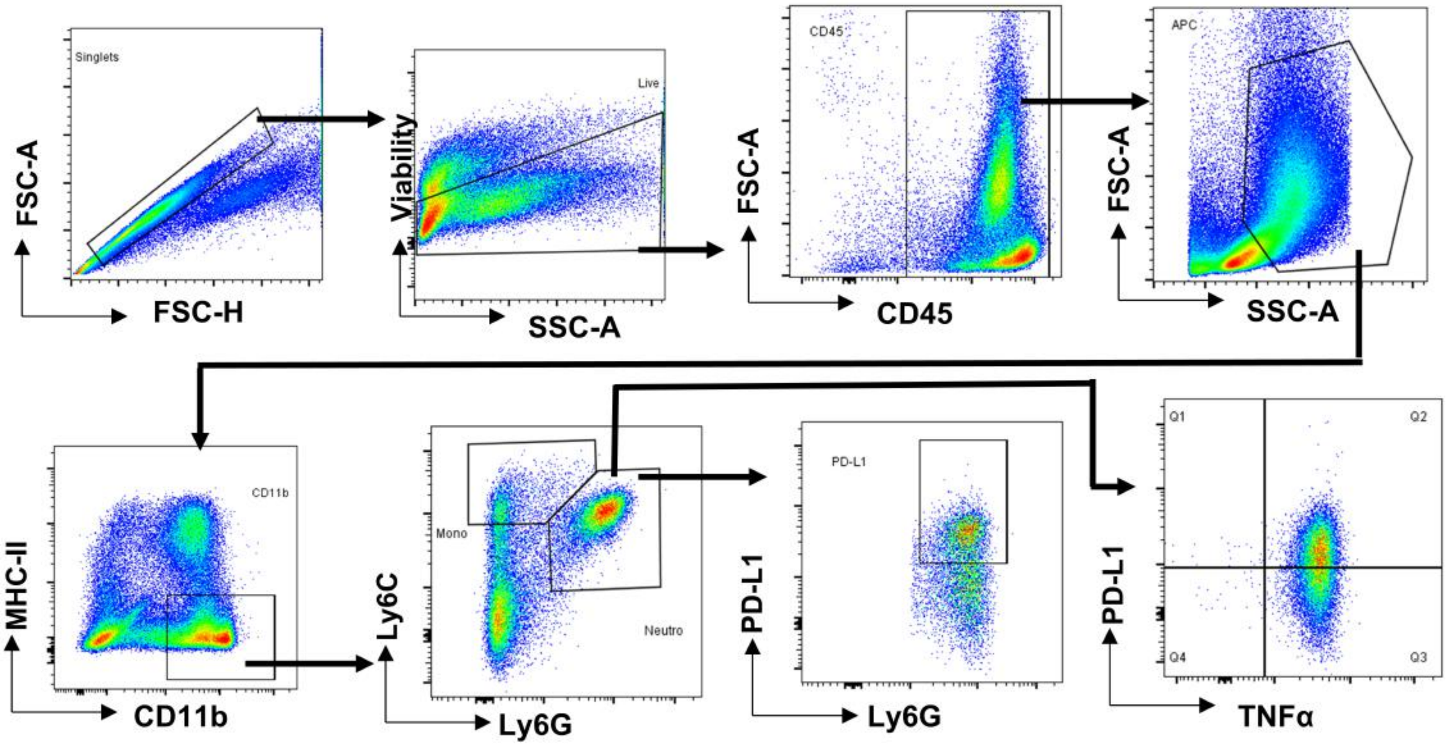
