## Supplementary Figure 8 for "PDL-1+ Neutrophils mediate susceptibility during endotoxemia in Metabolic Dysfunction–Associated Steatotic Liver Disease"

**Figure S8. TNFR1/R2 -/- animals not have reduced neutrophil infiltration in the hepatic tissue of animals with MAFLD during endotoxemia**

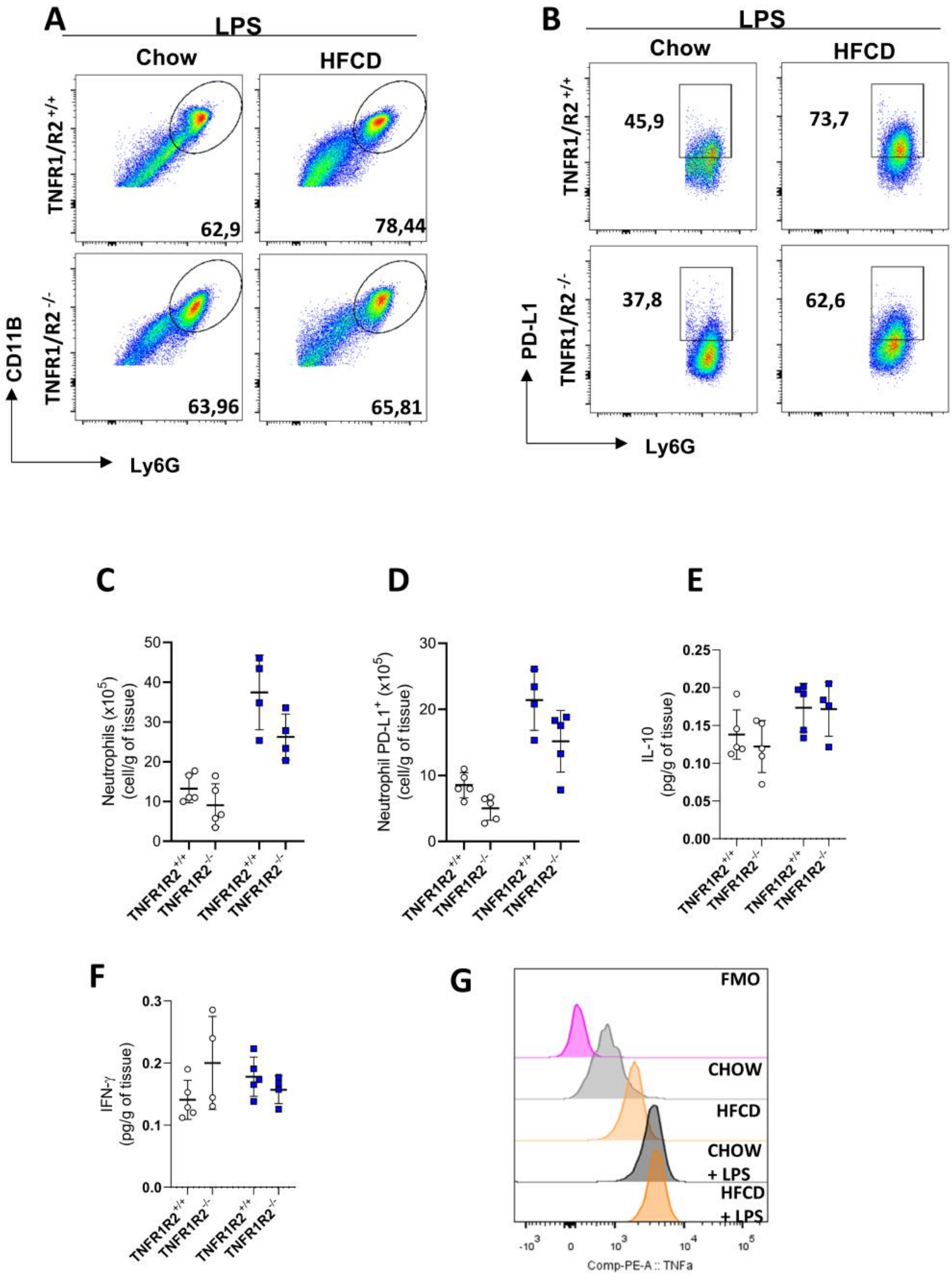
