## Supplementary Figure 9 for "PDL-1+ Neutrophils mediate susceptibility during endotoxemia in Metabolic Dysfunction–Associated Steatotic Liver Disease"

**Figure S9. PD-L1 expression in neutrophils occurs in the liver and not in the lung in animals with MAFLD during endotoxemia**

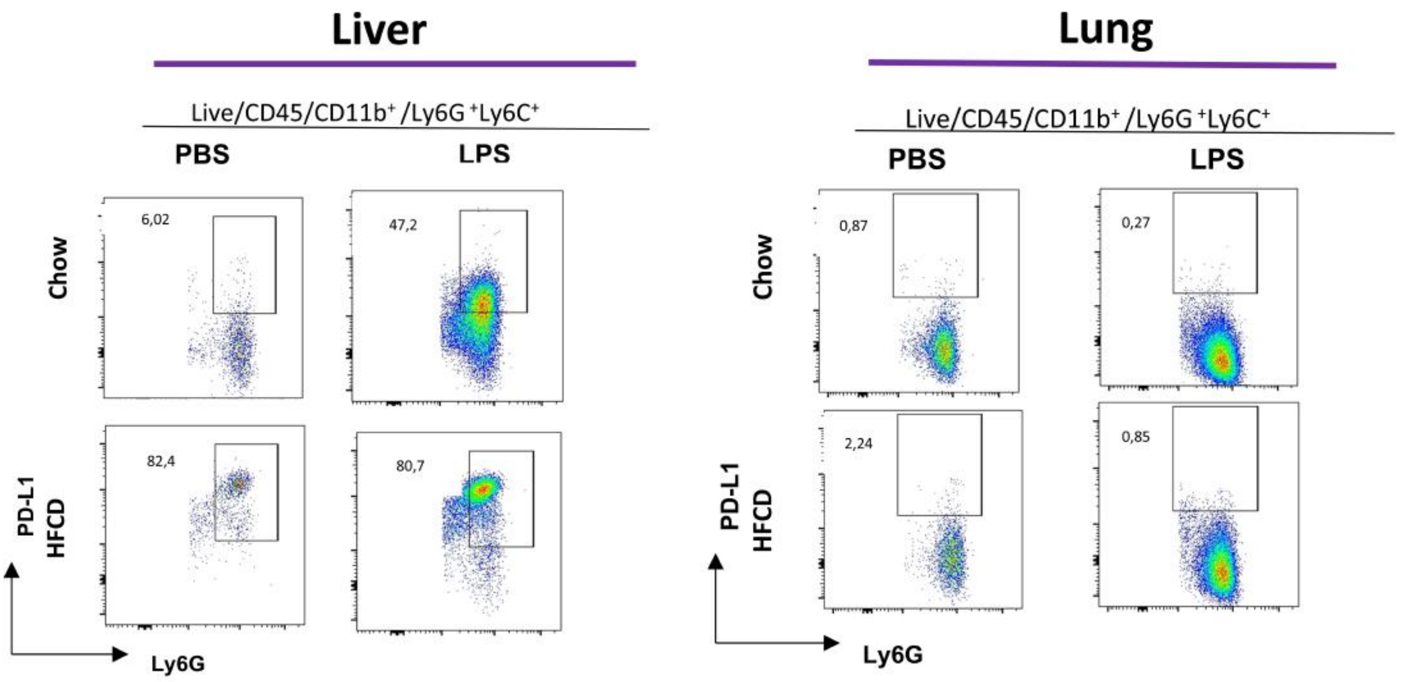
